# Human-supervised Agentic AI for Hypothesis Generation and Experimental Assistance in Drug Repurposing

**DOI:** 10.64898/2026.04.20.719538

**Authors:** Dinh Long Huynh, Elin Asp, Flavio Ballante, Jordi Carreras Puigvert, Alisa DeGrave, Reagon Karki, Kristen Nader, Päivi Östling, Bishab Pokharel, Jonne Rietdijk, Lars Schlotawa, Lina Schmidt, Srijit Seal, Brinton Seashore-Ludlow, Tero Aittokallio, Ola Spjuth

**Affiliations:** Department of Pharmaceutical Biosciences and Science for Life Laboratory, Uppsala University, Uppsala, Sweden; Department of Oncology and Pathology and Science for Life Laboratory, Karolinska Institutet, S-171 76 Stockholm, Sweden; Chemical Biology Consortium Sweden (CBCS), Science for Life Laboratory, Department of Medical Biochemistry and Biophysics, Karolinska Institutet, Tomtebodavägen 23A, 17165 Solna, Sweden; Pixl Bio AB, Uppsala, Sweden; Fraunhofer Institute for Translational Medicine and Pharmacology ITMP, Translational Neuroinflammation and Automated Microscopy TNM, 37083 Göttingen, Germany; Fraunhofer Institute for Translational Medicine and Pharmacology ITMP, Discovery Research Screening Port, Schnackenburgallee 114, 22525 Hamburg, Germany; Institute for Molecular Medicine Finland (FIMM), HiLIFE, University of Helsinki, Helsinki, Finland; Department of Paediatrics and Adolescent Medicine, University Medical Centre Göttingen (Universitätsmedizin Göttingen), Göttingen, Germany

## Abstract

Computational drug repurposing has focused largely on rapid hypothesis generation, yet real-world applications span a much broader lifecycle: suggesting drug candidates, designing experiments, analyzing assay data, and iteratively refining hypotheses. Here, we demonstrate that agentic AI can operate throughout this lifecycle. We developed RepurAgent, a hierarchical multi-agentic system comprising a supervisor agent and a planning agent that coordinates four specialized sub-agents (research, prediction, data, and report) with human in the loop, supported by episodic memory and retrieval-augmented generation. The system is grounded in repurposing-specific data, tools, and standard operating procedures developed within the REMEDi4ALL consortium. We validated RepurAgent across three scenarios spanning distinct stages within the repurposing lifecycle: in acute myeloid leukemia, a blinded expert evaluation indicated that RepurAgent produced substantially more candidates that were both novel and mechanistically credible than a vanilla LLM baseline; in a retrospective COVID-19 antiviral screen, RepurAgent acted as an adaptive experimental collaborator, prioritizing compounds with AUC-ROC up to 0.99 and average precision up to 0.91 without predefined thresholds and flagging confounders missed in manual review; and for multiple sulfatase deficiency, it prioritized 81 high-confidence candidates from 5159 compounds, which were further corroborated by domain experts. These results show that agentic AI can support work across the drug repurposing lifecycle, from hypothesis generation to experimental analysis. RepurAgent is open source and deployed at https://repuragent.serve.scilifelab.se/.

## Introduction

Drug repurposing, the systematic identification of new indications for existing approved drugs, has become a productive strategy in drug discovery, offering shorter development timelines and lower costs because it builds on established safety and pharmacokinetic profiles^1–5^. The COVID-19 pandemic showed this potential, with remdesivir and baricitinib rapidly repurposed and approved for antiviral and anti-inflammatory applications within months of the outbreak^6–9^. Beyond infectious disease, repurposing has produced new treatments in oncology, rare diseases, and neurology, including thalidomide for multiple myeloma^10,11^ and sildenafil for pulmonary arterial hypertension^12,13^. Each of these successes required connecting drug mechanisms, biological targets, and disease pathophysiology across many heterogeneous data sources. As the number of approved drugs and characterized diseases grows, doing this manually becomes increasingly impractical^2,14,15^, motivating the development of computational approaches.

Early computational approaches relied on chemical similarity^16,17^, guilt-by-association methods^18,19^, and network-based reasoning^20–22^ to identify candidate drug-disease associations. Knowledge graphs (KGs) then provided a principled framework for integrating heterogeneous biomedical entities, including drugs, targets, pathways, phenotypes, and diseases, into structured representations suited to computational reasoning. Prominent examples include Hetionet^14^, which integrated 29 public resources into a heterogeneous network, and BioKG^23^, which extended this to drug-specific relationships. Building on these representations, machine learning (ML) and deep learning (DL) methods have enabled relationship prediction at scale, with approaches ranging from graph neural networks and embedding models such as TransE^24–26^ and RotatE^27–29^, to transformer-based architectures trained on multi-modal biomedical data. More recently, large language models (LLMs) have been applied to extract and reason over biomedical literature^30–34^, which extends the evidence available to repurposing pipelines beyond structured databases.

Despite this progress, existing computational approaches remain fundamentally fragmented. KGs integrate structured data well but must be rebuilt manually as biomedical knowledge evolves; widely used KGs such as Hetionet^14^, DRKG^27^, and OREGANO^35^ last updated in 2016, 2020, and 2023 respectively. ML models for relationship prediction are powerful but narrowly scoped, requiring pre-processed, structured input and producing ranked candidate lists without mechanistic justification. Literature mining tools retrieve evidence without connecting it to predictive pipelines. Absorption, distribution, metabolism, excretion, and toxicity (ADMET) prediction, pathway enrichment, and report generation each require separate tools with incompatible input and output formats. Integrating these components and reasoning across them iteratively still falls mostly to the human scientist. Most of this effort lies in the steps between tools, including retrieving and harmonizing heterogeneous data, analyzing experimental results, detecting confounders, and reprioritizing candidate compounds as wet-lab results come in. Most computational frameworks address one of these steps in isolation, so coordinating them into a continuous workflow that spans both hypothesis generation and experimental support remains largely a manual task.

Agentic AI systems, in which LLMs are equipped with external tools, memory, and reasoning loops to plan and execute multi-step tasks^36–38^, could address this fragmentation problem. Unlike static ML pipelines, agents can dynamically select and invoke tools, adapt their reasoning in response to intermediate outputs, and maintain context across workflows^39–44^. Biomedical AI agents have been demonstrated across domains: ChemCrow^40^ augmented LLMs with chemistry tools for molecular design, CellAtria^42^ developed an agentic framework for single-cell RNA-seq analysis, and DeepRare^41^ applied traceable reasoning to rare disease diagnosis. Agents have also been developed for drug repurposing itself: DrugAgent^45^ coordinates ML model, knowledge graph, and literature-retrieval agents to assess drug-target interactions, and ChatDRex^46^ gives conversational access to network-based disease module identification and drug prioritization over the NeDRex knowledge graph. Both operate *in silico* on hypothesis generation, and neither covers the analysis of laboratory experimental data. General-purpose scientific agents have shown that a single system can span hypothesis generation and experimental analysis. The Google AI co-scientist^44^ generates and refines hypotheses through multi-agentic debate, and the repurposing candidates it proposed for acute myeloid leukemia were later tested *in vitro.* Robin^47^ couples literature-search agents with a data analysis agent to propose therapeutic hypotheses, interpret the results, and revise its hypotheses; it identified ripasudil as a candidate treatment for dry age-related macular degeneration. Kosmos^48^ uses a structured world model to coordinate data analysis and literature search over extended autonomous runs. These systems reason mainly through literature agents and general-purpose analysis code, and they are designed for high autonomy. The curated resources that repurposing relies on (disease-specific KGs, compound annotations, ADMET prediction) and the standard operating procedures (SOPs) that repurposing laboratories already follow are not central to their design.

An agentic system that combines these resources and SOPs with support for the experimental stages of a campaign has therefore received less attention. This gap matters in practice because a ranked candidate list only starts a repurposing campaign. As experimental results accumulate, the team has to retrieve, integrate, and act on heterogeneous evidence. Here, we present RepurAgent, a multi-agentic AI system developed to operate throughout the repurposing lifecycle, from candidate generation to the analysis of experimental results. We ground the system over data, tools, and SOPs for drug repurposing developed within the Horizon Europe REMEDi4ALL consortium, and evaluate it in three scenarios that together span different stages within the repurposing lifecycle: autonomous end-to-end candidate generation for acute myeloid leukemia, adaptive experimental collaboration in a COVID-19 antiviral screen, and hypothesis-driven prioritization for the ultra-rare disease multiple sulfatase deficiency. These case studies show how agentic orchestration grounded in repurposing-specific tools and SOPs can address not only the prediction problem but also the wider scientific workflow in which prediction is embedded.

## Results

### System overview

RepurAgent is a multi-agentic AI system designed to assist across multiple stages of drug repurposing, from hypothesis generation to experimental data analysis. Incoming requests are first routed to the *planning agent*, which produces a structured, multi-step workflow that human scientists can review and refine before execution (**Figure 1a**). Once the plan is approved, the supervisor agent coordinates the execution under human supervision. It delegates sub-tasks to four specialized sub-agents (a *research agent*, a *prediction agent*, a *data agent*, and a *report agent*), collects their outputs, routes to the next task, and adapts the remaining plan as needed.

**Figure 1.**
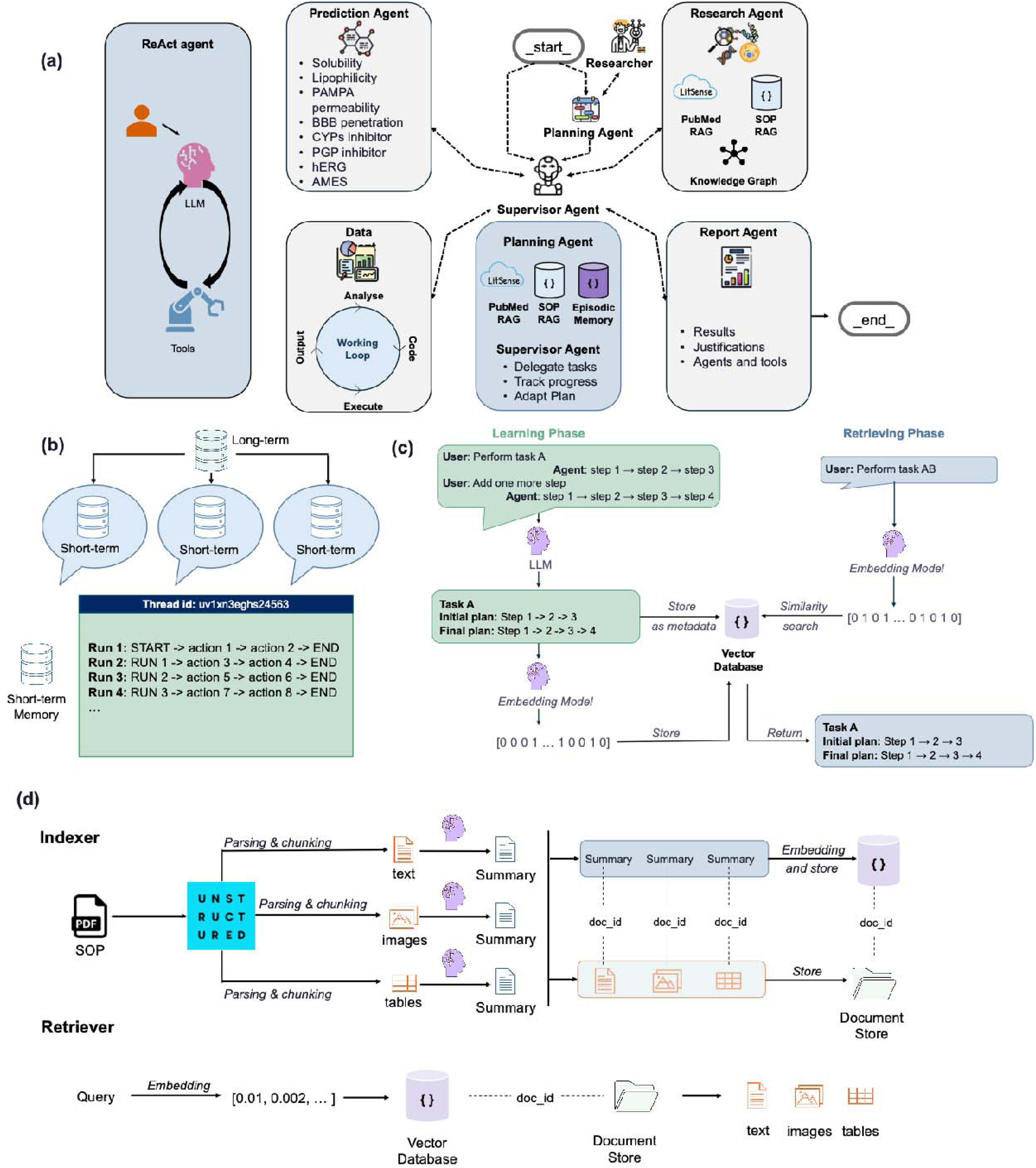
System architecture of RepurAgent. (a) Multi-agent architecture of RepurAgent. The planning agent and the supervisor agent coordinate the workflow: the plan is refined through multi-turn interaction with a human scientist and, once approved, sub-tasks are delegated to the specialized data, research, prediction, and report agents. (b) Memory architecture comprising short-term memory, retained within a single conversation, and long-term episodic memory, which persists across sessions. (c) Learning and retrieval phases of the episodic memory module. (d) Retrieval-augmented generation (RAG) framework incorporating REMEDi4ALL standard operating procedures (SOPs) as an internal knowledge base to support planning, reasoning, and decision-making.

Each agent operates using the ReAct^49^ pattern, combining specialized tools with domain-specific system prompts. The *research agent* retrieves and synthesizes biomedical knowledge through live API calls rather than static snapshots, so its information stays current as biomedical evidence evolves. Within the REMEDi4ALL consortium, multiple tools and resources for drug repurposing were developed or adapted for AI agent access: a knowledge graph generator (KGG)^50^ for real-time disease-specific KG construction; the RepurposeDrugs^51^ predictive platform; and the Chemical Annotator^52^, a unified annotation toolkit that combines data from ChEMBL^53^, PubChem^54^, UniChem^55^, and KEGG^56^ to retrieve physicochemical properties, molecular targets, and quantitative activity measurements for compounds. Consortium-validated SOPs for drug repurposing established within the REMEDi4ALL consortium were also incorporated to guide the agents’ reasoning (**Table S1**). The *prediction agent* supports ADMET property prediction using a suite of pre-trained conformal prediction models. The *data agent* autonomously generates and executes Python code to read experimental datasets, perform analysing algorithms, and inspect outputs iteratively, while the *report agent* synthesizes findings into structured, human-readable summaries. Together, these components equip RepurAgent with a comprehensive, domain-grounded toolkit that spans the full repurposing workflow: from real-time KG construction and live biomedical database retrieval, through mechanistic reasoning over drug targets and pathways, to property prediction and structured report generation. The full list of tools available to each agent is given in **Table S2**.

To support continuity across sessions, RepurAgent implements a three-layer memory architecture comprising working memory (graph state shared among agents during a session), short-term memory (conversation context persisting across turns within a session), and long-term episodic memory (a vector-indexed knowledge base of task decompositions that the user chose to save and will be retrieved semantically to inform future planning; **Supplementary Note 1**, **Figure 1b-c).** Retrieval-augmented generation (RAG) over internal SOPs further grounds the system in consortium-specific experimental protocols. At runtime, the retriever performs semantic search to surface the most relevant guidance into the agent’s context (**Figure 1d**). This keeps agent reasoning aligned with established laboratory practice.

### Planning agent performance is maximized by tool augmentation and requires human oversight

To assess how much each of the planning agent’s tools contributes to plan quality, we evaluated 8 configurations (**Table S3**) across 15 drug repurposing task requests, including 5 requests from the real case studies and 10 synthetic requests covering other repurposing scenarios. The full request set, system prompt, and evaluation protocol, together with per-run judge scores, are available in a Zenodo repository (**Data Availability**). Each request was evaluated over 10 runs with an LLM-as-judge framework that scores plan quality against a set of criteria and rubrics (**Table S4**), and scores were normalized to a 0-1 scale. A Kruskal-Wallis test confirmed significant variation in quality across configurations (*p* < 10^-100^), and pairwise Mann-Whitney U tests with Holm correction showed that every tool-augmented configuration significantly outperformed the vanilla LLM baseline (all adjusted *p* < 10^-10^; **Figure 2a-b**).

**Figure 2.**
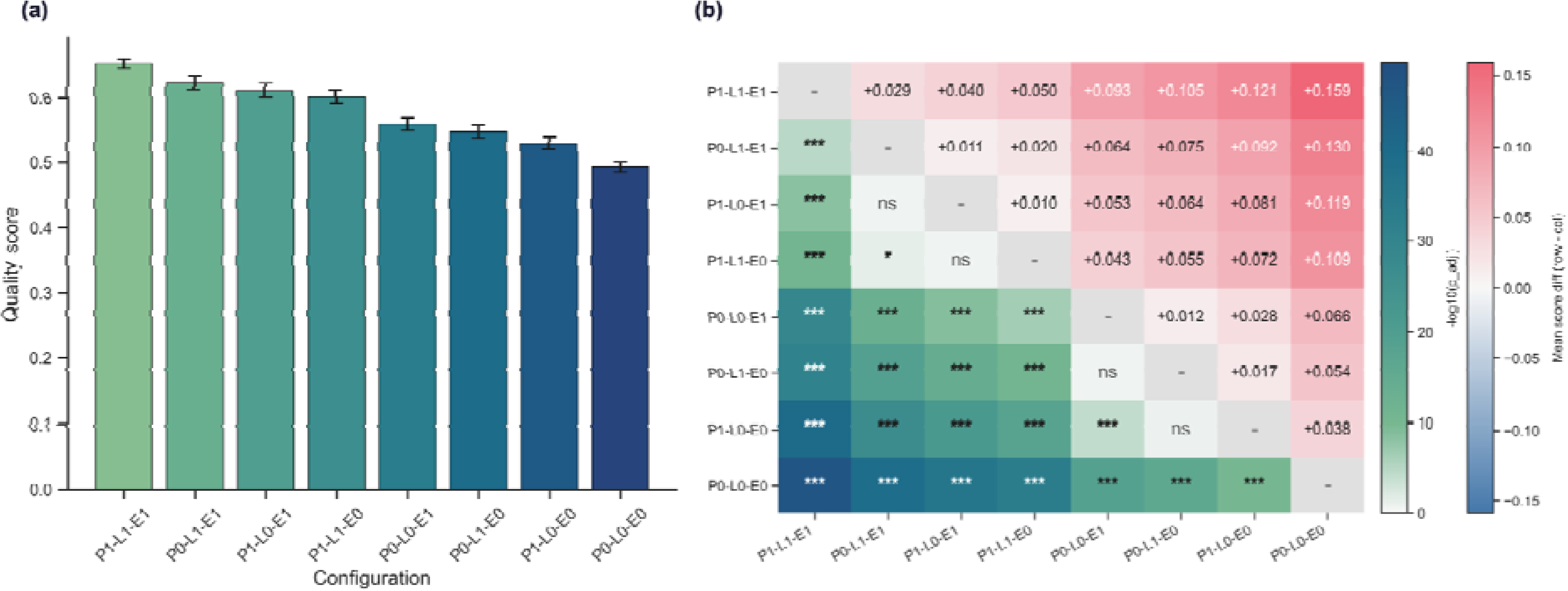
Ablation study for the planning agent. (a) Bar chart showing the mean quality score for each evaluated configuration, ranked in descending order. Error bars represent 95% confidence interval (CI). Higher scores indicate plans rated as higher quality by the LLM judge. (b) Pairwise comparison heatmap across all configurations. Lower triangle (green scale): statistical significance of pairwise Mann-Whitney U tests, expressed as −log10(p-adjusted); darker shading indicates stronger evidence of a difference in plan quality. Upper triangle (pink-blue scale): mean score difference between row and column configurations; pink indicates superior performance of the row configuration, blue indicates the opposite. All p-values were corrected for multiple comparisons using the Holm method. Significance thresholds: *** p < 0.001, ** p < 0.01, * p < 0.05; ns, not significant (p ≥ 0.05). Configuration labels follow the notation P{0,1}-L{0,1}-E{0,1}, where P, L, and E denote the presence (1) or absence (0) of standard operating procedures, literature retrieval, and episodic memory, respectively. All p-values were corrected for multiple comparisons using the Holm method; *** p < 0.001, ** p < 0.01, * p < 0.05; ns, not significant (p ≥ 0.05).

The baseline vanilla LLM (P0-L0-E0) achieved a composite quality score of 0.492. Each tool individually provided modest but significant improvements: SOP retrieval alone (P1-L0-E0) raised the score by 0.038, PubMed literature search alone (P0-L1-E0) by 0.054, and episodic memory alone (P0-L0-E1) by 0.066 (**Figure 2a-b**). The fully augmented configuration (P1-L1-E1) achieved the highest composite score of 0.651, a 32% improvement over the baseline (**Figure 2a-b**).

Even so, the best configuration reached only a quality score of 0.651, indicating that autonomous planning alone cannot yet decompose complex scientific tasks reliably. Scientists should therefore review, correct, and refine agent-generated plans before execution, to ensure that the system’s reasoning aligns with domain expertise and study-specific constraints (**Figure 1a**).

### RepurAgent executes an end-to-end workflow to identify repurposing candidates for acute myeloid leukemia

Acute myeloid leukemia (AML) is an aggressive hematological malignancy driven by recurrent somatic mutations, including in FLT3, NPM1, DNMT3A, and IDH1/2, which disrupt differentiation, promote aberrant proliferation, and sustain leukemic stem cell populations^57–59^. Despite high remission rates with standard induction chemotherapy^60^, relapse rates remain high due to clonal evolution and therapy-resistant subclones, which makes AML an important disease for drug repurposing.

Starting from a single natural language prompt, RepurAgent orchestrated a repurposing workflow from knowledge acquisition to candidate prioritization, without predefined pipelines or external datasets. The scientist refined the plan during the planning phase to adjust the operation strategy (**Table S5**), and human supervision is recommended throughout execution to monitor intermediate outputs. During this process, the system constructed an AML-specific KG integrating known drugs, protein targets, mechanisms of action, pathways, and side-effect information from Open Targets^61^, GWAS^62^, ChEMBL^53^, and UniProt^63^. The resulting KG captured how disease-relevant biological entities connect, so that downstream analysis could prioritize biologically plausible repurposing candidates. More details of this workflow are described in **Figure 3a**. Top-ranked candidates targeted a range of proteins, including FMS-like tyrosine kinases, platelet-derived growth factor receptors, protein kinase C-related kinases, ribosomal proteins, and DNA polymerases (**Figure S1**). Among these targets, FLT3, the most commonly mutated kinase in AML, is directly targeted by four suggested agents (sunitinib, dovitinib, famitinib, and fedratinib). Together, these top 20 candidates implicated key AML-relevant pathways including PI3K/AKT, MAPK/ERK, JAK/STAT, RUNX1-regulated hematopoietic differentiation, cell-cycle regulation, and apoptosis (**Figure S2-3**). Beyond the top 20, BCL2-targeting agents (obatoclax, navitoclax) and IDH1/2 inhibitors (vorasidenib, enasidenib) also appeared in the ranked list, but at ranks 119-171 of the 397 compounds in the initial drug space.

**Figure 3.**
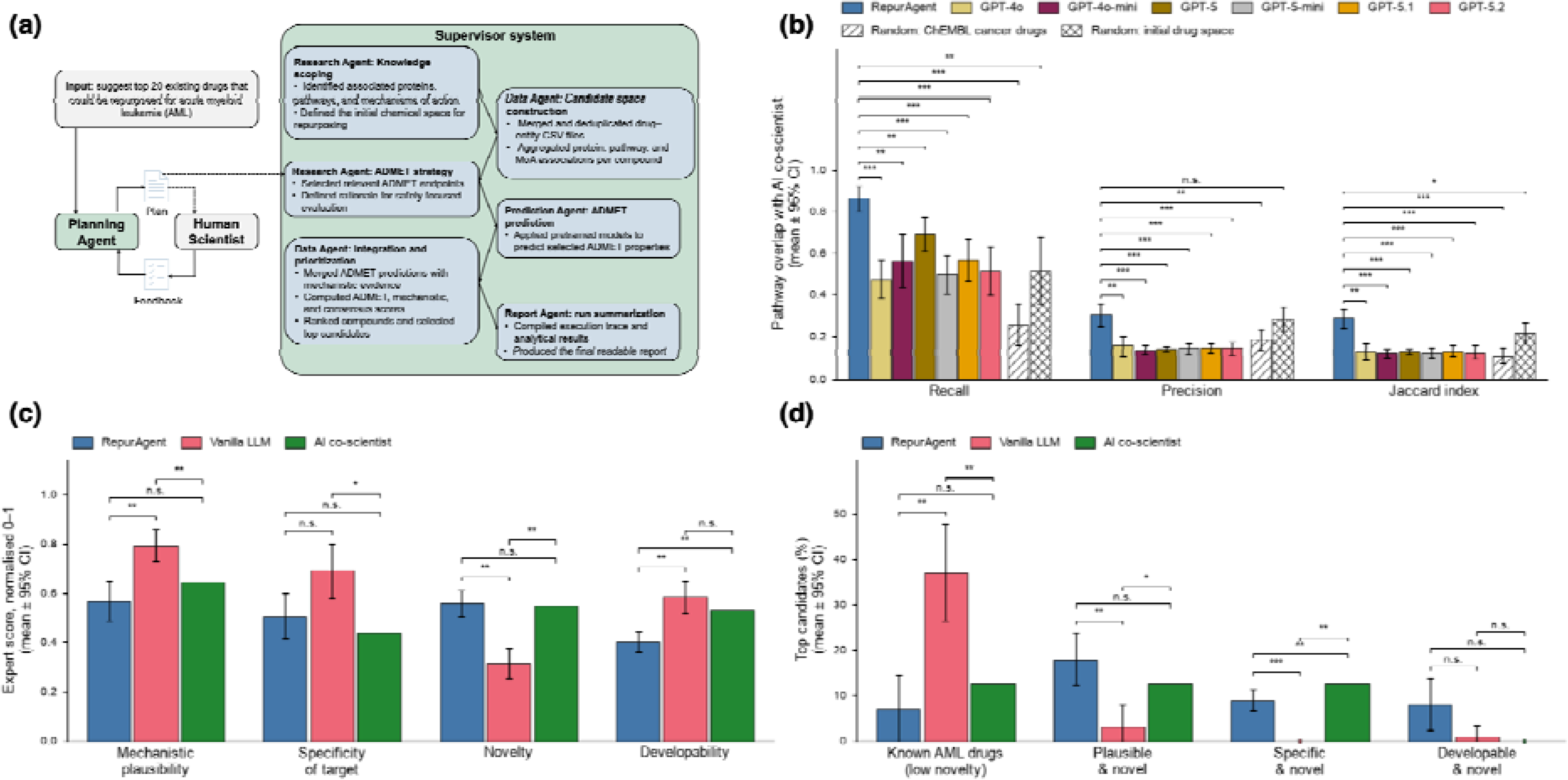
Benchmarking RepurAgent against vanilla LLMs and Google Co-Scientist in the AML case study. (a) RepurAgent workflow for identifying drug repurposing candidates for acute myeloid leukaemia (AML). A human scientist supervises the planning phase together with the planning agent. Once the scientist approves the plan, the supervisor agent executes it end to end, coordinating the specialised agents for knowledge scoping, ADMET strategy definition, candidate space construction, ADMET prediction, data integration and prioritisation, and report generation. (b) Recall, precision, and Jaccard index of pathway overlap with Google Co-Scientist’s candidates (reference set) for RepurAgent, six vanilla LLMs, and two random baselines: drugs randomized from 3941 ChEMBL cancer drugs, and drugs randomized from RepurAgent’s 397-compound initial drug space. Bars show the mean across runs, and error bars are 95% confidence intervals. Brackets compare each arm with RepurAgent by two-sided Mann–Whitney U test, with Holm correction across all 24 comparisons. RepurAgent had the highest overlap on all three metrics (recall 0.86, precision 0.30, Jaccard index 0.28) and differed significantly from every arm, except for precision against random drugs from its own initial drug space (0.28; p = 0.68). (c) Expert scores on four rubric criteria, normalized to a 0-1 scale. Bars show the mean across runs and error bars are 95% confidence intervals across runs. Brackets show Holm-adjusted p-values for the three pairwise comparisons within each criterion: RepurAgent vs vanilla LLM by two-sided Mann– Whitney U test, and each of these systems vs Google Co-Scientist by two-sided one-sample Wilcoxon signed-rank test with the Co-Scientist score as a fixed reference (**Methods**). (d) Percentage of each system’s candidates in four derived categories. Percentages were computed per run, and bars show their mean; a dot on the axis marks 0%. Error bars, tests and Holm correction are as in (c). In all panels: n.s., p ≥ 0.05; *p < 0.05; **p < 0.01; ***p < 0.001.

We benchmarked RepurAgent against vanilla LLMs and the Google Co-Scientist pipeline^64^. The Co-Scientist workflow is expert-intensive, integrating agent-generated scores with DepMap data and oncology expert review at multiple stages before arriving at five final repurposing candidates (**Figure S4**), spanning the RAS-RAF-MEK-ERK, JAK-STAT, FLT3, Keap1-Nrf2, NFκB, and Mevalonate pathways. Because Google Co-Scientist published only its five final drug candidates without disclosing the initial drug library or intermediate selection steps, a direct drug-level comparison is not feasible. We therefore compared pathway coverage as a proxy, which captures the mechanistic space covered rather than the specific drugs selected. RepurAgent’s top 20 candidates covered 86% of the pathways associated with Co-Scientist’s selected drugs (**Figure 3b, Table S6**). This was significantly higher than all six vanilla LLMs on recall, precision, and Jaccard index.

To test whether the RepurAgent’s pathway coverage simply reflects generic or pan-cancer pathways that any oncology drug set would recover, rather than AML-relevant biology, we added a random cancer-drug baseline: from a pool of 3941 cancer-associated drugs in ChEMBL, we drew 10 independent random samples of 20 drugs each and ran the same pathway enrichment and overlap analysis on them (**Figure 3b**). This baseline recovered only 26% of Co-Scientist’s pathways, well below RepurAgent’s 86%. RepurAgent significantly outperformed the random cancer-drug baseline on recall, precision, and Jaccard index (all Holm-adjusted p < 0.01; **Figure 3b**). Because a random cancer-drug set would itself recover any generic or pan-cancer pathways, its low overlap indicates that RepurAgent’s convergence with Co-Scientist reflects selection of AML-relevant pathways rather than an artifact of broad oncology pathway coverage. A second question is how much of this overlap is already determined by the initial drug space that RepurAgent assembled from its AML KG, before any prioritization. We therefore added a second random baseline, drawing 10 random samples of 20 compounds from the 397 compounds in RepurAgent’s initial drug space. These random draws recovered 52% of Co-Scientist’s pathways, twice the random cancer-drug baseline but well below RepurAgent’s prioritized top 20 (86%; Holm-adjusted p = 0.009), and reached a lower Jaccard index (0.22 vs 0.28; p = 0.037). Their precision, however, was comparable to RepurAgent’s (0.28 vs 0.30; p = 0.68). This suggests that the KG-derived drug space is itself enriched for Co-Scientist-relevant pathways and largely accounts for RepurAgent’s precision advantage, while the prioritization step adds pathway coverage.

In a sensitivity analysis that removed broad Reactome terms (pathways with gene sets larger than N genes, with N = 500, 300, 200, 100, 50, and 20) from the pathway sets, RepurAgent remained above the random cancer-drug baseline and the vanilla LLMs on recall, precision, and Jaccard index at all N thresholds (Holm-adjusted p ≤ 0.016; **Figure S5**, **Table S7**).

Additionally, we ran a blinded expert evaluation of the top-ranked drugs from RepurAgent, a vanilla LLM (GPT-5.2), and Google Co-Scientist for the same AML task. Deduplicating the top candidates across the 3 systems and 10 runs yielded 63 unique compounds. Each was scored on two independent evaluation forms (126 drug-form evaluations). The experts saw only the drug name, primary indication, and development phase and were blinded to the reasoning behind each drug and to the system that proposed it. Candidates were rated 1-5 scale and normalized to 0-1 scale on mechanistic plausibility (A1), target/pathway specificity (A2), novelty of the drug-AML hypothesis (A3), and druggability/developability (A4). The detailed criteria and rubrics were described in the **Methods** section and in **Table S8**. Scores were averaged within each run, and systems were compared on these run-level scores (**Methods**). Compared with RepurAgent, the vanilla LLM scored higher on mechanistic plausibility (0.79 vs 0.57; p-value = 0.002), target specificity (0.69 vs 0.51; p-value = 0.05), and developability (0.58 vs 0.40; p-value = 0.004, **Figure 3c**). RepurAgent scored higher on novelty (0.56 vs 0.31; p-value = 0.002, **Figure 3c**). Because Co-Scientist has only a single candidate set and we could not re-run it externally, we compared the 10 runs of each other system (RepurAgent and vanilla LLM) against Co-Scientist scores as a fixed reference (0.64, 0.44, 0.55 and 0.53 for A1–A4). The vanilla LLM runs scored above Co-Scientist on plausibility (p = 0.008) and specificity (p = 0.012) and below it on novelty in all 10 runs (p = 0.004). RepurAgent did not differ from Co-Scientist on plausibility, specificity or novelty (p = 0.08, 0.13 and 0.82), but all 10 of its runs scored below Co-Scientist on developability (p = 0.004). Further analysis revealed that the vanilla LLM’s high scores for mechanistic plausibility and target specificity reflect a bias toward drugs already established in AML rather than genuine repurposing: on average 37% of vanilla LLM’s top candidates were known AML drugs (novelty score ≤ 2; e.g. gilteritinib, quizartinib, midostaurin, azacitidine), compared with 7% of RepurAgent (p = 0.004) and 12.5% for Co-Scientist (p = 0.008; **Figure 3d**). Because such drugs are approved for AML, they score high on plausibility, specificity and developability while contributing nothing new. Therefore, we quantified the fraction of candidates that were simultaneously novel and biologically relevant (**Figure 3d**). On this basis, RepurAgent outperformed the vanilla LLM and was indistinguishable from Co-Scientist on two of the three composite categories. RepurAgent proposed more plausible-and-novel (18% vs 3%; p = 0.003) and specific-and-novel candidates (9% vs 0%; p < 0.001) than the vanilla LLM, while the difference in developable-and-novel candidates was not significant (8% vs 1%; p = 0.07). Co-Scientist (12.5%, 12.5% and 0%) also exceeded the vanilla LLM on the first two categories (p = 0.012 and p = 0.004) and did not differ from RepurAgent on plausible-and-novel (p = 0.07) or developable-and-novel candidates (p = 0.07). These results indicated that RepurAgent trades some plausibility for substantially more novel yet mechanistically credible hypotheses than the vanilla LLM, with a profile closer to that of the expert-intensive Co-Scientist workflow.

### RepurAgent as an active assistant for COVID-19 repurposing screening project

In a retrospective analysis, we applied RepurAgent to the completed experimental dataset from Asp et al.^65^ to evaluate whether the agent could reproduce and extend expert analytical decisions across the three-stage screening workflow within a fraction of the time. The original study used host-centric targets for COVID-19 drug repurposing, in which researchers ran three primary screens of a 5275-compound library in Vero E6 cells, namely a cytopathic effect inhibition assay, an antibody-based screen, and a Cell Painting assay. This yielded 324 primary hits, defined as compounds active in at least one assay. The team then performed additional cell painting assay, antibody-based and phospholipidosis screens on human A549-ACE2 cells to narrow down the list to 74 hits, to retain compounds with antiviral effects and remove those active only because of phospholipidosis, which is a membrane accumulation artifact known to produce false positives in antiviral assays^66^. Finally, the researchers manually evaluated the targets, mechanisms of action, and physicochemical features of these 74 hits and selected five potential candidates for further investigation (**Figure 4a**). Manual data analysis for this study required several weeks to complete.

**Figure 4.**
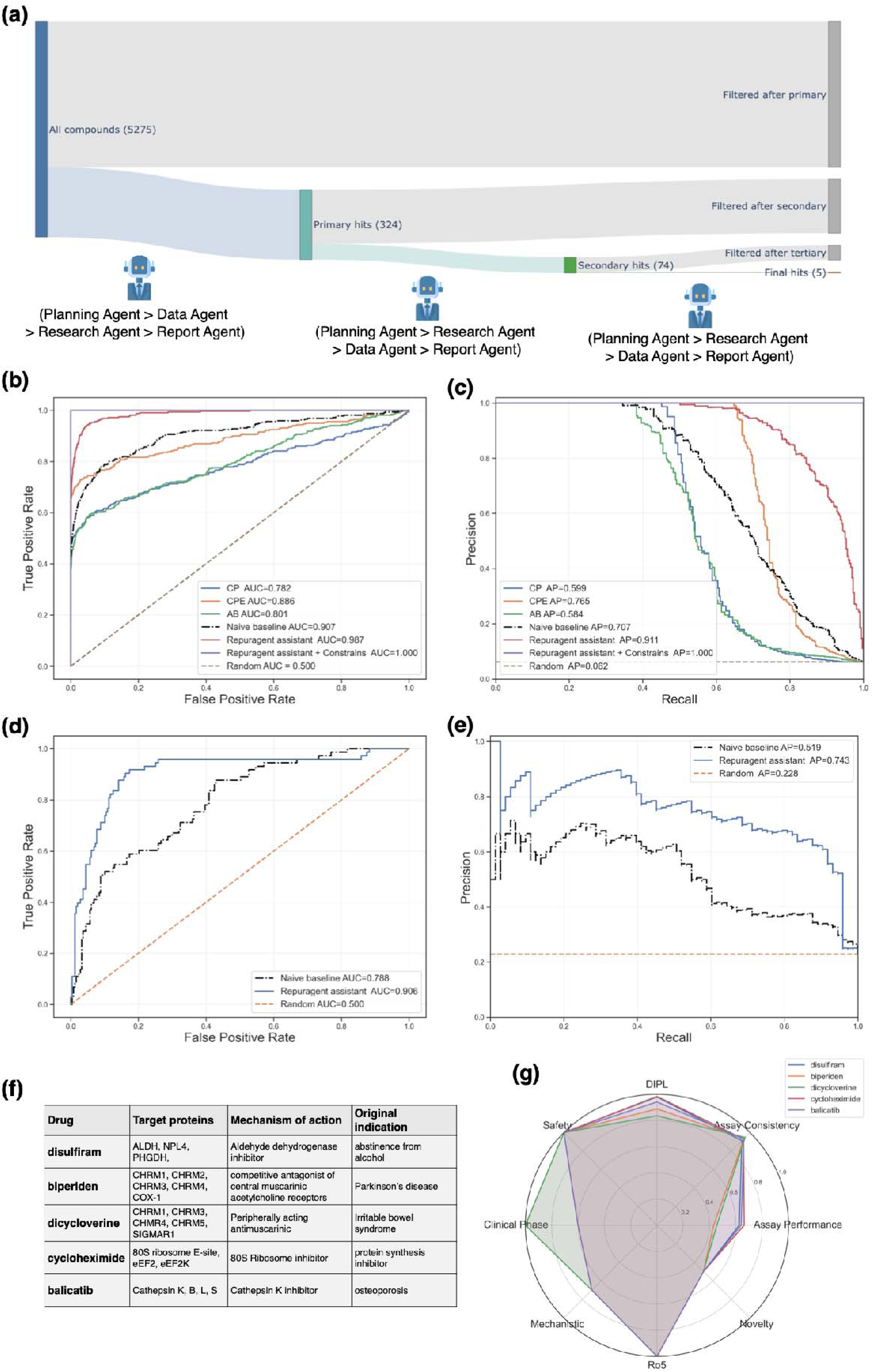
RepurAgent-assisted analysis of a COVID-19 drug repurposing screen. (a) Experimental workflow adapted from Asp et al.^65^, comprising a primary screening phase in Vero E6 cells (324 hits), followed by validation assays in human A549 cells (74 hits) and final manual review to select five candidates; RepurAgent assists at each stage with data analysis, hypothesis monitoring, confounder identification, and experimental recommendations. (b, c) Primary screen performance: ROC (b) and precision-recall (c) curves for ranking the 5275 compounds against the 324 expert-selected hits. Curves are shown for RepurAgent with the published activity thresholds supplied (constrained) and without them (unconstrained), for a fixed mean-aggregation baseline, and for each single assay readout: Cell Painting (CP) morphology score, cytopathic effect (CPE) inhibition and antibody-based (AB) infection rate. (d, e) Validation screen performance: ROC (d) and precision-recall (e) curves for ranking the 324 re-screened primary hits against the 74 expert-selected secondary hits, for RepurAgent and the naive baseline. Legends report the area under the curve (AUC) in ROC panels and the average precision (AP) in precision-recall panels. (f) Top five candidate drugs proposed by RepurAgent based on integrated analysis of all experimental and externally retrieved data. (g) Radar plot depicting the multi-criteria evaluation profile and comparison of the five selected repurposing candidates.

RepurAgent was employed to assist this iterative workflow (**Figure 4a**). Each screening stage was run as a separate session, and the scientist revised the plan before approval in each (**Table S5**). Given the raw readouts from the three primary assays and the approved plan, RepurAgent autonomously merged, analyzed, and ranked the compounds by antiviral potency. The agent can do this in two modes. When the scientist can specify the desired activity thresholds, the agent applies them faithfully (constrained mode); when such criteria are not available, the agent infers a ranking on its own (unconstrained mode). We first tested the constrained mode by supplying the agent with the activity thresholds published by Asp et al.^65^ as a transparent rule-based reference. Under these predefined rules, the agent recovered the expert hit list exactly (AUC-ROC = 1.00) (**Figure 4b**). In the unconstrained mode, the agent inferred the thresholds itself and achieved an AUC-ROC of 0.987 (**Figure 4b**). Because the dataset is highly imbalanced (only 6.2% of the primary library were hits), we also report the precision-recall curve alongside the ROC curve. In the unconstrained mode, RepurAgent reached an AP of 0.911 (**Figure 4c**), and 81.6% of its top 324 compounds were expert-selected hits (**Table S9).** The self-derived ranking therefore recovered the expert manual selection accurately without predefined rules. To reach this ranking, the agent integrated orthogonal readouts from cell-painting morphology, antibody-based infection, and cytopathic effect (CPE) rescue assays through normalization and weighted aggregation. The procedure remains fully transparent, allowing scientists to rerun the analysis with custom weightings if desired. Beyond ranking, RepurAgent also detected potential confounders by examining inconsistencies across the experimental readouts. For example, it flagged compounds that showed potent infection inhibition in the antibody-based assay while reducing host cell viability by more than 20%, indicating that the apparent antiviral effect may instead reflect cytotoxicity. A complete list of confounder warnings is provided in **Table S10**. RepurAgent then analyzed the validation screen for the 324 primary hits in A549-ACE2 cells to assess performance and detect further confounders. The agent achieved an AUC-ROC of 0.906 and an AP of 0.743 (**Figure 4d, e),** and 69.9% of its top 74 compounds were expert-selected hits (**Table S9**). Here, the agent also identified further confounders: cases in which cell-painting morphological rescue occurred without a corresponding reduction in infection (**Table S10**).

To isolate the contribution of the agent’s reasoning from that of simple data aggregation, we introduced a fixed, non-agentic comparator at both screening stages, which is simply taking means of the same normalized readouts, ranked without any agentic reasoning or tool use (**Methods**). On the primary screen, this baseline reached an AUC-ROC of 0.907 and an AP of 0.707, compared with AUC 0.987 and AP 0.911 for RepurAgent; on the validation screen, it reached AUC 0.788 and AP 0.519, compared with AUC 0.906 and AP 0.743 of RepurAgent (**Figure 4b-e**). At a cutoff equal to the number of expert-selected hits, 64.2% and 54.8% of the baseline’s selections were hits, compared with 81.6% and 69.9% for RepurAgent (**Table S9**). These results indicate that the agent’s adaptive weighting improves on static aggregation at both screening stages, most clearly among the top-ranked compounds that would be carried forward. The baseline also still requires an analyst to build the pipeline, including parsing, harmonizing, and normalizing the raw assay files and specifying the aggregation scheme, before any score can be computed, while RepurAgent performed these steps autonomously from the raw experimental outputs. For this task, therefore, the benefit of the agent also lies in its orchestration of data cleaning and integration across both screening stages.

The final 74 hits were reranked using the data from both the primary and validation screens. RepurAgent employed a ranking algorithm that considered multiple criteria, including assay performance, consistency across assays, drug-likeness, mechanistic support, safety profile, and clinical trial status. After completing the analysis, RepurAgent generated a structured report for each of the top five drugs: disulfiram, biperiden, dicycloverine, cycloheximide, and balicatib. **Figure 4f** lists the biological target, mechanism of action (MoA), and original indication of each drug, and **Figure 4g** shows why each was selected. Expert review confirmed that these five drugs could have potential for pandemic preparedness. Three of them (disulfiram, biperiden, and dicycloverine) are approved oral agents, which would facilitate rapid deployment. Overall, RepurAgent reproduced the expert ranking decisions in minutes, compared with several weeks of manual analysis, at comparable analytical quality.

### RepurAgent prioritises promising drugs to repurpose for Multiple Sulfatase Deficiency

In this case study, RepurAgent was tasked with prioritizing 5159 compounds for multiple sulfatase deficiency (MSD) in terms of their repurposing potential and classifying them into high, medium, and low confidence tiers. MSD is an ultra-rare disease caused by pathogenic variants in the *SUMF1* gene, which encodes the Formylglycine-Generating Enzyme (FGE)^67,68^. Mutated FGE impairs the post-translational activation of all 17 sulfatases, leading to lysosomal accumulation of glycosaminoglycans and sulfolipids^69–71^ and convergent downstream pathologies including neurodegeneration^72^, astrocyte dysfunction^73^, and neuroinflammation^74^.

The main challenge in this case study was the extreme scarcity of annotated disease-specific data: the initial MSD KG generated by RepurAgent contained only 46 nodes, anchored to four genes (SUMF1, SUMF2, ARSA, ARSB) (**Figure 5a**) whose direct pharmacological targeting is either non-specific or beyond the scope of drug repurposing. To compensate, RepurAgent supplemented KG with PubMed literature retrieval and synthesized the evidence to define three mechanistic therapeutic directions: FGE chaperone stabilization, lysosomal rescue, and symptomatic inflammatory rescue. These directions were operationalized into a three-tier ranking framework, which external domain experts independently validated as a comprehensive representation of MSD pathophysiology. As in the other case studies, the scientist revised the plan before approval; in this session the revision only merged steps, and the therapeutic directions and confidence-tier criteria were derived by the agent (**Table S5**).

**Figure 5.**
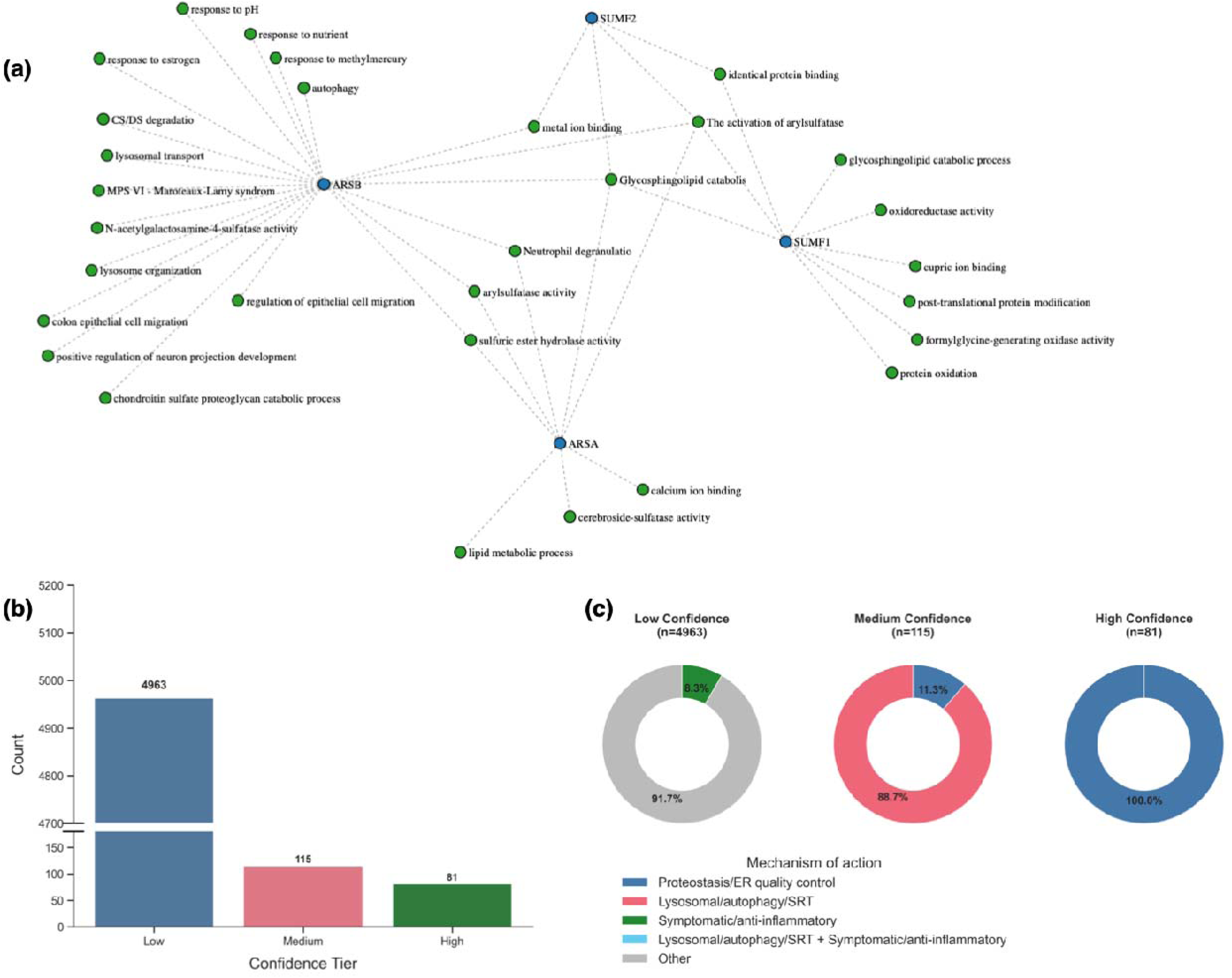
RepurAgent prioritized compounds for the ultra-rare disease, multiple sulfatase deficiency (MSD). (a) MSD-specific KG describing the disease biological landscape. Nodes representing gene products are shown in blue (ARSA, ARSB, SUMF1, SUMF2) and associated GO terms in green, with edges indicating annotated functional relationships. The KG highlights shared pathways including lysosomal function, glycosphingolipid catabolism, sulfatase activity, neutrophil degranulation, and autophagy, reflecting the multifaceted pathophysiology of MSD. (b) Number of compounds in each confidence tier assigned by RepurAgent. (c) Donut charts displaying the breakdown of MoA categories within each confidence tier. Low-confidence compounds are predominantly classified as "Other" (91.7%), with a minor fraction attributed to symptomatic control (8.3%). Medium-confidence compounds show a marked shift toward lysosomal/autophagy/SRT mechanisms (88.7%), with smaller contributions from proteostasis/ER control and combined MoA categories. High-confidence compounds are entirely attributed to proteostasis/ER control (100%).

To test whether the workflow extends to rare and data-poor indications, we applied RepurAgent across the 5159-compound library. The framework was highly selective: only 81 compounds were classified as high confidence and 115 as medium confidence (**Figure 5b**). High-confidence compounds were mechanistically homogeneous, targeting the proteostasis and endoplasmic reticulum quality control axis, while medium-confidence compounds were more diverse, with 88.7% mapped to lysosomal/autophagy/substrate reduction mechanisms (**Figure 5c**). Across confidence tiers, drug-target binding affinities formed a sparse interaction profile that was structured by mechanism category (**Figure S6**). Notably, external experts independently flagged one high-confidence compound (within the top 1.6%) as particularly promising on the basis of their own unpublished experimental findings, which corroborates RepurAgent’s prioritization.

## Discussion

Most computational tools for drug repurposing remain anchored to a single stage of the repurposing pipeline and require substantial manual effort to connect. The persistent challenges, thus, arise not within individual prediction steps but around them: retrieving and harmonizing heterogeneous data, analysing experimental results, detecting confounders, and reasoning over intermediate findings. These challenges are inherently dynamic, as the appropriate action at any given point depends on the evolving state of the analysis. The principal contribution of this work is therefore an architectural demonstration: AI agent orchestration, grounded in repurposing-specific tools and SOPs developed by experts from an international consortium, and supervised by the scientist, can coordinate the different stages of drug repurposing, spanning from hypothesis generation, candidate prioritization, to experimental data analysis, within one adaptive framework.

The three case studies illustrate complementary aspects of these capabilities. In the AML case, RepurAgent constructed a disease-specific KG and ranked candidates whose pathway coverage converged with that of the Google Co-Scientist candidates, which had been selected through an expert-intensive workflow. The two approaches arrive at largely overlapping pathway space (FLT/PDGFR, PI3K/AKT/mTOR, JAK/STAT, DNA repair, **Table S6**) through fundamentally different processes. This suggests that systematic database traversal guided by agentic reasoning can recover much of the mechanistic space that expert review arrived at, at least at the level of pathway coverage. The two random baselines indicate where this convergence arises: random draws from cancer drugs had significantly lower coverage, while random draws from RepurAgent’s KG-derived drug space had higher coverage than the cancer-drug draws but still lower than RepurAgent (**Figure 3b**), suggesting that both KG construction and prioritization contribute. By contrast, vanilla LLMs suggested drugs already approved or in late-stage trials for AML, which reflects their reliance on parametric recall of frequently co-occurring drug-disease pairs. The blinded expert evaluation quantified this with 37% of the vanilla LLM’s candidates were known AML drugs, compared with 7% for RepurAgent, which explains why the vanilla LLM scored higher on plausibility and developability but lower on novelty (**Figure 3c, d**). Grounding candidate generation in live database queries reduced this bias, at a cost: RepurAgent’s candidates scored lower on average plausibility, but more of them were both plausible and novel than those of the vanilla LLM (18% vs 3%). Against the single published Co-Scientist candidate set, RepurAgent’s runs did not differ consistently in 6 of 8 comparisons. The COVID-19 case study shifts the focus from hypothesis generation to experimental assistance: against a fixed equal-weight aggregation of the same readouts, the agent’s adaptive weighting improved the ranking at both screening stages (AUC-ROC 0.987 vs 0.907, AP 0.911 vs 0.707 for primary screen and AUC 0.906 vs 0.788, AP 0.743 vs 0.519 for validation screen). However, concordance is measured against expert-curated selections, not a prospective biological endpoint; the intended contribution of this case study is assistance with the analytical workflow, including autonomous parsing, harmonisation, normalisation, and aggregation of heterogeneous raw assay files, together with confounder detection. Because the fixed baselines still require a data analyst to build the pipeline before any score is produced, the orchestration of these steps is where the agent removes analyst effort. The MSD case study tests the opposite data regime: when the disease-specific KG contained only 46 nodes and four genes, the agent compensated by extending its reasoning with literature retrieval and organizing the disease into three therapeutic directions, a strategy independently validated by domain experts. Together, these cases show that agentic AI can operate across the range of data availability found in real-world repurposing campaigns.

The COVID-19 case study also shows how SOPs can act as practical guardrails for agentic reasoning. SOPs contributed during planning, where their inclusion improved plan quality (**Figure 2**), and during the analysis itself, where they supplied specific decision rules. At the validation screen stage, retrieval of two phospholipidosis-specific SOPs provided the threshold that compounds inducing cell counts below 80% of DMSO controls should be flagged as toxic, a rule the agent applied directly. At the final selection stage, a retrieved *in silico* repurposing SOP instructed prioritization of compounds that have reached the market (Molecule Max Phase = 4), which RepurAgent operationalized as a clinical readiness sub-score, while physicochemical guidance on logP was translated into penalties for rule-of-five violations and extreme lipophilicity. These are not post hoc interpretations; they form a retrievable chain from SOP content through the agent’s reasoning to specific parameter choices. More broadly, SOPs act as soft-constraint policies: they steer reasoning toward established best practices without hard-coding agent behavior. In experimental drug repurposing, the validity of an analysis often turns on procedural knowledge: toxicity thresholds, assay weighting conventions, and confounder definitions that language models are unlikely to have absorbed from training data alone but that formalized SOPs routinely encode. Systematic collection, curation, and open publication of SOPs, as undertaken by the REMEDi4ALL consortium, therefore supports the reliability of AI-assisted scientific workflows.

A central concern for AI agents is whether their outputs, beyond being plausible, remain inspectable and grounded in verifiable evidence. RepurAgent addresses this in four ways: a human approval step precedes plan execution (**Table S5**); tool-using agents are instructed to rely strictly on retrieved evidence and avoid fabricated assertions; tools provide structured failure responses rather than failing silently; and the Python execution environment is constrained through import allowlists and restricted function access to prevent unintended side effects. RepurAgent also exposes its full execution trace to the scientist, rather than deliver only the final output. This transparency is particularly important in long-running workflows, where silent error propagation can compromise downstream conclusions. Other agentic systems share this emphasis: DeepRare^41^ grounds its reasoning in traceable, verifiable references; CellAtria^42^ emphasizes transparency and auditability as design objectives; and ChemCrow^40^ demonstrates why expert assessment remains indispensable when factual precision matters.

Other agentic systems for therapeutic discovery fall into two groups: repurposing-specific and general-purpose agents. The former shares RepurAgent’s reliance on structured evidence: DrugAgent^45^ reconciles machine learning predictions, knowledge graph paths, and literature evidence for individual drug-target pairs, and ChatDRex^46^ orchestrates established network-medicine algorithms over the NeDRex knowledge graph, with a reasoning module that keeps the user in the loop. Both focus on *in silico* hypothesis generation. RepurAgent applies a comparable tool-grounded design to experimental screening data as well, as shown in the COVID-19 case study. General-purpose scientific agents go further than RepurAgent in prospective validation and in scale: Robin^47^ tested its hypotheses in the laboratory, and ripasudil was confirmed *in vitro* as a candidate for dry age-related macular degeneration; our COVID-19 analysis, by comparison, is retrospective. Kosmos^48^ uses a structured world model to keep its agents coherent across about 200 rollouts, a scale of autonomous investigation that RepurAgent does not attempt. Both systems are built for high autonomy, and Kosmos, for example, does not yet let scientists intervene between cycles. RepurAgent instead routes its reasoning through curated repurposing resources and consortium SOPs, and the scientist must approve an explicit plan before anything is executed. We therefore regard RepurAgent as complementary to these systems.

This study has several limitations. Prospective wet-lab validation remains the most important next step: the AML candidates are experimentally untested, the corroborated MSD candidate cannot be disclosed due to intellectual property constraints, and the COVID-19 study is retrospective. The AML expert evaluation has further constraints that Google Co-Scientist is an external system that we could not re-run, so it was evaluated on its single published candidate set as a fixed reference. This comparison ignored both Co-Scientist’s run-to-run variance, so differences from Co-Scientist describe whether a system’s runs were consistently above or below this one output, not a general difference between systems. In addition, the three case studies evaluate fundamentally different forms of utility and cannot be reduced to a single performance metric. This reflects the diversity of real-world repurposing scenarios and complicates systematic benchmarking. More broadly, the field currently lacks community-accepted benchmarks for end-to-end agentic systems, and constructing them is complicated by the fact that literature-aware agents may retrieve published answers rather than infer them independently, especially for benchmarks derived retrospectively from validated drug-disease associations. In this study, we addressed memorization by design in two of the three case studies. In the AML case study, the vanilla LLM baseline served as a memorization-only arm: it favored drugs already established in AML and achieved lower pathway overlap with Co-Scientist than RepurAgent, whose top-ranked candidates all originated from knowledge-graph tool outputs. In the MSD case study, the corroborating evidence consists of the experts’ unpublished experimental findings, which cannot be present in any training corpus. The COVID-19 case study re-analyzes a published screen, so we relied on an audit of the agent’s traces. The audit showed that the rankings came from agent-written scoring rules that contained no compound-specific selections. However, this audit cannot rule out that prior knowledge shaped the agent’s analytical choices, particularly because the knowledge cutoff of GPT-5.2 (the LLM behind the data agent) is August 2025, which postdates the first public reports of the screening study^75^. A future systematic evaluation of memorization will therefore require benchmarks built on data released after model training or held privately, combined with controlled tests such as masking of compound identities. Finally, the contribution of human supervision was assessed descriptively. In every case-study session the scientist revised the plan before approval (**Table S5**), but the raw unrevised plans were not executed, so the effect of these revisions on the final outputs relative to fully autonomous execution was not quantified.

Because RepurAgent is modular, its capabilities are not fixed at deployment. New tools, such as additional database integrations, alternative KG sources, and new ADMET predictors, can be incorporated without restructuring the agent framework, and new SOPs can be indexed into the retrieval system to expand domain coverage. The system therefore becomes more useful as domain-grounded tools and protocols that anchor its reasoning in established scientific practice accumulate. As prospective validation links computation and experiment, the central question will shift from whether AI agents can support drug repurposing to how the community can equip them to do so reliably and responsibly.

In summary, we show that agentic AI can operate across connected stages of drug repurposing, supporting experimental workflows as well as hypothesis generation. RepurAgent, grounded in data, tools, and consortium-validated SOPs, coordinates four specialized sub-agents through a human-supervised reasoning loop. In three case studies, covering a hematological malignancy, an infectious-disease screening campaign, and an ultra-rare disease with sparse biological annotation, RepurAgent achieved competitive mechanistic coverage, high concordance with expert compound selection, and autonomous confounder detection. The principal advance is the orchestration layer that connects evidence retrieval, mechanistic reasoning, property prediction, and experimental data analysis into a single continuous and auditable workflow. As agentic AI systems mature and prospective wet-lab validation closes the loop between computation and experiment, human-supervised agentic orchestration may become a practical component of routine drug repurposing workflows.

## Methods

### AI Agent framework

RepurAgent is implemented in Python and built on LangChain (v0.3.24) and LangGraph (v0.6.3). The system is deployed as a multi-user web application with Gradio (v5.49.1). The workflow follows a tool-augmented ReAct pattern: at each step, a LLM produces either a natural-language response or a structured tool call; tool outputs are appended to the message state and used in subsequent reasoning steps. The LLM for each agent was chosen according to the agent’s role and the results of iterative system-prompt tuning (**Table 1**).

**Table 1.** LLM used by each agent.

| **Component** | **LLMs** | **Release date** | **Knowledge cutoff date** |
|---|---|---|---|
| Planning agent | Gpt-4o | May 13, 2024 | Oct 01, 2023 |
| Supervisor agent | Gpt-5-mini | Aug 7, 2025 | May 31, 2024 |
| Research agent | Gpt-5-mini | Aug 7, 2025 | May 31, 2024 |
| Data Agent | Gpt-5.2 | Dec 11, 2025 | Aug 31, 2025 |
| Prediction Agent | Gpt-5-mini | Aug 7, 2025 | May 31, 2024 |
| Report Agent | Gpt-5-mini | Aug 7, 2025 | May 31, 2024 |

A memory architecture was adopted to support both within-session and cross-session context. Short-term memory was implemented using LangChain’s PostgreSQL checkpointer (v2.0.3), which persists the full conversation state across turns within an active session. Episodic memory was managed through ChromaDB (v0.6.3) in conjunction with LangMem (v0.2.9), enabling retrieval-augmented recall of relevant context across sessions (more details in **Supplementary Note 1**).

SOP documents were ingested through a multi-stage preprocessing pipeline. Raw PDF files were first parsed using Unstructured (v0.18.4), which extracted text, images, and tabular content as discrete components. Each component was subsequently chunked and enriched with a natural-language description generated by GPT-5.1, providing semantic context for modalities that lack inherent textual representation, such as figures and tables. Vector representations were then computed using the text-embedding-3-small embedding model and indexed in the vector store for downstream retrieval.

### Standard Operating Procedures (SOPs) for drug repurposing

Within the REMEDi4ALL consortium, a range of SOPs for drug repurposing has been developed and published in a Zenodo community (https://zenodo.org/communities/remedi4all) (**Table S1**). Work to extend the list is ongoing and will further inform the reasoning of RepurAgent and of other AI agents for drug repurposing that use this resource.

### Ablation study of the planning agent

Beyond the base LLM, the planning agent was augmented with three additional tools: protocol_search_sop, literature_search_pubmed, and episodic_memory. Enabling or disabling each tool produced a 2^3 factorial design of 8 configurations, denoted P{0,1}-L{0,1}-E{0,1} for SOP (P), literature (L), and episodic memory (E) retrieval (**Table S3**).

We constructed a benchmark of 15 planning requests, each paired with an expert-defined reference decomposition. Five requests were derived from the three real case studies (AML, COVID-19, and MSD), and the remaining ten were synthetic requests spanning a range of repurposing scenarios, including disease-landscape synthesis, host-directed antiviral review, ADMET and safety triage, oral-bioavailability and CNS prioritisation, phospholipidosis triage, mechanism and pathway clustering, and combination pre-selection. Reference decompositions ranged from two to six steps. The complete task set and reference decompositions are publicly available in the Zenodo repository (**Data Availability**).

For each configuration, the planning agent received a request together with the retrieved context available under that condition and returned a structured plan comprising a goal, an ordered list of steps, an objective, an expected output, and a note on critical success factors. Each plan was scored by an independent LLM-as-judge model (GPT-4o), instructed to act as a strict reviewer. The judge was provided with the request, the reference decomposition, the set of enabled tools, the retrieved context, and the generated plan, and rated the plan across six dimensions: structure and completeness, task alignment, executability, scientific rigour, context use, and specificity. Each dimension was scored on a 1 to 5 scale with rubric anchors defined at levels 1, 3, and 5 (**Table S4**). Scores were normalized to the 0 to 1 interval, and the composite quality score was taken as the mean across the six dimensions. To account for stochastic variation, each configuration was evaluated over ten independent runs per task, yielding 1200 generated plans in total (15 tasks × 8 configurations × 10 runs).

Differences in scores across configurations were evaluated with a two-stage non-parametric procedure. First, a Kruskal-Wallis H-test assessed whether any systematic difference existed across all configurations. Where it indicated significant variation, pairwise two-sided Mann-Whitney U tests compared every pair of configurations. Raw p-values were adjusted using the Holm correction. Differences were considered significant at *p*-value < 0.05.

### Evaluation of RAG performance

To assess retrieval performance, we constructed a stratified benchmark of 200 queries relevant to the SOPs, spanning diverse query types. Retrieval quality was evaluated at two levels: file level and evidence level, where the correct source document must appear within the retrieved set and where the exact supporting passage must be retrieved, respectively. Performance was quantified using two metrics.

Recall@k measures the proportion of queries for which at least one relevant item appears within the top-*k* retrieved results:

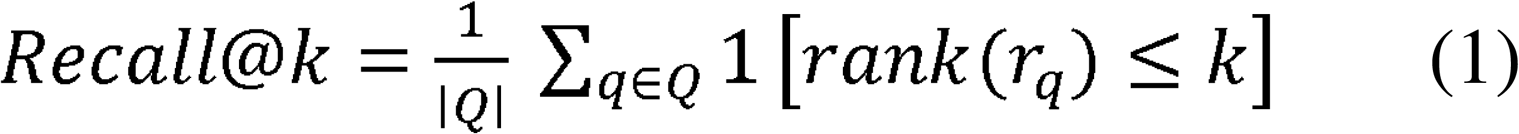

where Q is the set of all queries, *r_q_* denotes the ground-truth relevant item for query *q*, and 1[·] is the indicator function. All evaluations used k = 4, consistent with the default retrieval depth of the system.

Mean Reciprocal Rank (MRR) captures the system’s ability to rank the most relevant result highly, by averaging the reciprocal of the rank at which the first relevant item appears:

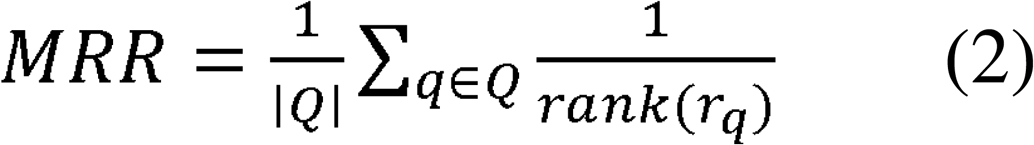

Where *rank*(*r_q_*) is the position of the first relevant item in the ranked list for query *q*. If no relevant item is retrieved, the reciprocal rank is set to zero.

Overall, the retriever demonstrated strong file-level performance, achieving a Recall@4 of 0.950 and a Mean Reciprocal Rank (MRR) of 0.883, indicating that the correct source document was identified within the top four results for the large majority of queries and was typically ranked near the top. Evidence-level performance was lower but remained robust, with Recall@4 = 0.795 and MRR = 0.607 (**Figure S7a-b**), reflecting the inherently greater difficulty of localizing precise text spans.

Performance varied systematically across query types (**Figure S7c-d**). Several categories achieved perfect or near-perfect file-level Recall@4 (1.0), including goal lookup, quality control, selection rule, filter rule, database lookup, reagent lookup, parameter lookup, equipment lookup, safety lookup, and toxicity rule. At the evidence level, filter rule and database lookup queries also reached Recall@4 = 1.0, while parameter lookup and selection rule queries achieved 1.0 and 0.929, respectively (**Figure S7c**). Evidence-level MRR was highest for quality control (0.787), metric lookup (0.777), and selection rule (0.696), reflecting strong result prioritization for these categories.

In contrast, software lookup queries presented the greatest challenge, yielding the lowest evidence-level Recall@4 (0.375) and MRR (0.292), likely due to semantic variability or sparse representation of software-related content in the indexed corpus. Equipment lookup (evidence-level Recall@4 = 0.667, MRR = 0.475), safety lookup (0.583, 0.559), and toxicity rule (0.667, 0.600) also underperformed relative to the overall average, identifying targeted categories for future improvement. Taken together, these results demonstrate that the retriever reliably identifies correct source documents across diverse query types while maintaining reasonable evidence-level precision, with specific improvement opportunities in software-, equipment-, and safety-related categories.

### ADMET prediction models

ADMET prediction is performed by calling pretrained CPSign (v2.0.0)^76^ models. Pre-trained CPSign models are provided for multiple endpoints, including CYP inhibition classifiers (CYP1A2/2C9/2C19/2D6/3A4), hERG, AMES, P-glycoprotein (PGP), PAMPA permeability, BBB penetration, and a solubility regressor. Training data were downloaded from Therapeutic Data Commons^77^ (TDC). The models were trained and hyperparameter-optimized, and predictions were precomputed, with the CPSign package. The model validation results are summarized in **Table S11**. CPSign employs conformal prediction, in which a confidence level defines the proportion of prediction sets that contain the true label. Classification models could return a single label, a multi-label set, or an empty set depending on the specified confidence. A confidence level of 0.80 was applied to all classification models and 0.71 to the solubility regressor. The exception is lipophilicity, which is computed directly from SMILES using RDKit’s Crippen logP^78^ because this parameter is often used in the literature and in guidelines.

### Pathways enrichment analysis in AML case study

To determine drug-pathway associations in the AML case study, pathway enrichment analysis was performed for each drug individually. Molecular targets for each compound were retrieved from the ChEMBL^53^ database and used as gene set inputs for enrichment analysis. Over-representation analysis against the Reactome_2022^79^ human gene set library was then conducted via Enrichr^80^, accessed programmatically through the GSEApy^81^ package (v1.1.11). Enrichment was performed independently for each drug, and pathways with a Benjamini-Hochberg adjusted *p*-value < 0.05 were considered statistically significant.

### Evaluation of pathways overlap with Co-Scientist in the AML case study

To assess how far each model’s repurposing candidates converge on the same biological pathway space as those of Google Co-Scientist, we computed three retrieval metrics (recall, precision, and Jaccard index), treating the Co-Scientist output as the reference set. Let denote the set of pathways returned by a Co-Scientist drug’s candidates and denote the set of pathways returned by a given model/agent’s candidates.

*Recall* is defined as the fraction of Co-Scientist pathways recovered by the model:

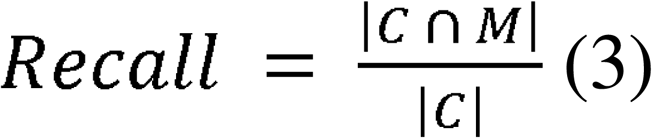

*Precision* is defined as the fraction of a model’s pathways that overlap with the Co-Scientist reference:

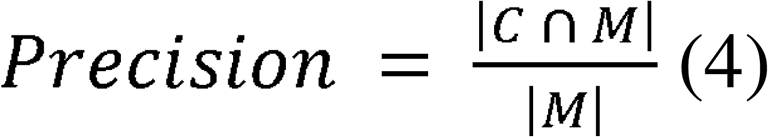

*Jaccard index* provides a symmetric measure of overall set similarity that jointly penalizes both missed and spurious pathways:

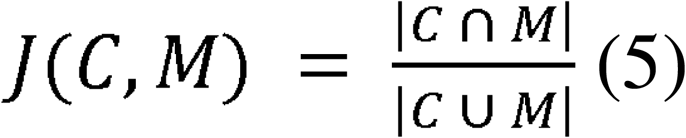

To establish a null expectation, we used two random baselines, each consisting of 10 independent random samples of 20 compounds (matching the size of RepurAgent’s top-20 list). Both baselines used the same random seeds, and every sample was processed through the same enrichment and overlap pipeline applied to RepurAgent’s candidates. The first baseline tests whether the overlap with Co-Scientist could arise from generic oncology pathways. For this baseline, we sampled from 3941 cancer-associated drugs collected from ChEMBL release 37 (accessed 13 July 2026 via the REST API). These were drawn from the ‘drug_indication’ endpoint, filtered to oncology indications, and restricted to compounds with a minimum phase ≥ 0.5. The set spans all anticancer indications and clinical phases, including both AML and non-AML agents. The second baseline separates the contribution of candidate-space construction from that of prioritization. For this baseline, we sampled from RepurAgent’s initial drug space: the 397 unique compounds it retrieved from the AML KG and associated biological entities.

All three metrics were computed per run across 10 independent runs for RepurAgent, each of the six vanilla LLMs, and each of the two random baselines. For each metric, group-level differences across the nine arms were first assessed using the Kruskal-Wallis test. Pairwise comparisons between RepurAgent and the eight arms (6 vanilla LLMs and 2 random arms) were then performed using two-sided Mann-Whitney U tests, and p-values were adjusted across all 24 pairwise comparisons (3 metrics × 8 arms) using the Holm-Bonferroni correction. Statistical significance was reported at three thresholds (*p ≤ 0.05, **p ≤ 0.01, ***p ≤ 0.001).

To assess the influence of broad pathway terms, we conducted a sensitivity analysis in which Reactome pathways with gene sets larger than N genes (N = 500, 300, 200, 100, 50, and 20) were progressively removed from both the Co-Scientist reference set and each arm’s pathway set. We then recomputed each metric and compared them using the same statistical tests as in the main analysis (**Figure S5**, **Table S7**).

### Expert evaluation in AML case study

#### Evaluation dataset preparation

We executed 10 independent runs of RepurAgent and 10 independent runs of a vanilla LLM baseline (GPT-5.2) with an identical task prompt, and retained the top-10 candidates from each run. These were combined with the eight candidates reported by Google Co-Scientist for the same task. Deduplicating across systems and runs yielded 63 unique drugs. A master provenance table was then set up to link each drug to its system and runs; this table was not shared with the experts conducting the evaluation.

### Expert panel and scoring rubric

Five evaluation forms were sent to experts in clinical and preclinical AML and haematological oncology. Each drug was assigned to two independent forms, giving 126 drug-form evaluations and 25-26 drugs per form. Candidates were shuffled before assignment, and experts saw only the drug name, its primary indication and its highest development phase. Experts were instructed not to attempt to infer the source and to rate each candidate on its own merits. Each candidate was scored on four criteria using a 1-5 scale: (A1) mechanistic plausibility for AML, (A2) specificity of the target or pathway to AML, (A3) novelty of the drug-AML hypothesis, and (A4) druggability and developability in the AML setting. A1 was explicitly framed as an assessment of the mechanism rather than of prior AML testing of that specific drug, so that an untested drug with well-established anti-leukaemic pharmacology could still score highly. The full rubric is given in **Supplementary Table S8**. Experts could also provide an optional free-text justification naming the strongest supporting argument and the principal weakness (A5).

### Score aggregation and unit of analysis

Completed forms were unblinded against the provenance table, and each score was propagated back to its original system and run. For each candidate and criterion, the experts’ ratings were averaged into a consensus score, and consensus scores were min-max scaled to the 0-1 range. For each system, run, and criterion, we averaged the consensus scores of the run’s candidates into a run-level score. This gave 10 run-level scores each for RepurAgent and the vanilla LLM and a single run-level score for Co-Scientist, for each criterion. All tests below operate on these run-level scores.

### Derived candidate categories

To distinguish established AML drugs from genuinely repurposable candidates, we classified each drug into four categories: known AML drugs (A3 ≤ 2, low novelty), plausible-and-novel (A1 > 3 and A3 > 3), specific-and-novel (A2 > 3 and A3 > 3), and developable- and-novel (A4 > 3 and A3 > 3). For each run, we computed the percentage of candidates meeting each category and analysed these percentages with the same procedures as the criterion scores.

### Comparison between RepurAgent and the vanilla LLM

With 10 run-level scores per system, we compared RepurAgent and the vanilla LLM using a two-sided Mann-Whitney U test. The test asks whether a RepurAgent run is more likely to score above or below a vanilla LLM run, without assuming that the scores are normally distributed.

### Comparison between RepurAgent/Vanilla LLM with the Co-Scientist

Co-Scientist has only one fixed candidate set, therefore its run-to-run variability is unidentifiable. To address this issue, we treated its run-level score as a fixed reference value. For each criterion, we computed the differences between each of the 10 run-level scores of RepurAgent (or the vanilla LLM) and the Co-Scientist score, and tested whether these differences are centered on zero with a two-sided one-sample Wilcoxon signed-rank test. This test assumes the Co-Scientist score as known exactly with no run-to-run variability. A significant result therefore indicates that a system’s runs scored consistently above or below this particular Co-Scientist output; it does not establish that the two systems differ in general.

Each criterion and each derived category was treated as a separate question, addressed by three pairwise comparisons (RepurAgent vs vanilla LLM, RepurAgent vs Co-Scientist, and vanilla LLM vs Co-Scientist). Therefore, we applied Holm’s correction across the three pairwise comparisons, with a two-sided significance level of 0.05. Significance marks in the figures and all p-values in the text are Holm-adjusted. The full results table, including Holm-adjusted p-values across all comparisons of each analysis, and the analysis code are available in the Zenodo repository (**Data Availability**).

### Benchmarking RepurAgent’s compound prioritization against expert selection in the COVID19 case study

To quantify the concordance between RepurAgent’s autonomous compound ranking and human expert selection in the COVID19 case study, we used receiver operating characteristic (ROC) and precision-recall analysis at two stages of the screening: the primary screen (5275 compounds; 324 expert-selected hits) and the validation screen (324 compounds; 74 expert-selected hits).

### Primary screen evaluation

RepurAgent received raw readouts (Excel files) from three orthogonal assays performed on Vero E6 cells: a Cell Painting (CP) morphological profiling assay, a cytopathic effect (CPE) inhibition assay, and an antibody-based (AB) infection assay. The agent autonomously merged and normalized these readouts, then produced a single ranked list of all 5275 compounds. We evaluated ranking quality under two conditions: (i) a constrained setting, in which the activity thresholds published by Asp et al.^65^ were supplied to the agent, and (ii) an unconstrained setting, in which RepurAgent inferred thresholds on its own. As single-assay baselines, we generated analogous ROC curves by sorting compounds on each individual readout: CP morphology score, CPE inhibition, and AB infection rate.

### Validation screen evaluation

For the 324 primary hits re-screened in human A549-ACE2 cells, RepurAgent independently ingested the validation-stage assay data and produced a new ranking. ROC analysis was performed against the 74 expert-selected hits using the rank-based procedure described below.

### ROC computation

For a given ranking or score, compounds were sorted and a classification threshold was swept across the ranked list. At each threshold, all compounds on the favorable side of the threshold were treated as predicted positives, and the true positive rate (TPR) and false positive rate (FPR) were computed as:

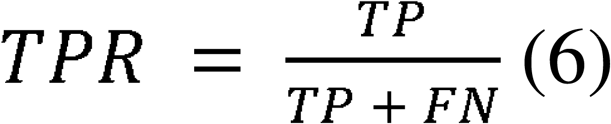

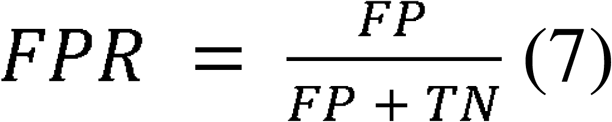

where TP is the number of expert-selected hits correctly classified as predicted positive (true positives), FN is the number of expert-selected hits not classified as predicted positives (false negatives), FP is the number of non-hits classified as predicted positive (false positives), and TN is the number of non-hits not classified as predicted positive (true negatives). The area under the ROC curve (AUC-ROC) was calculated by trapezoidal integration using sklearn.metrics.auc^82^. A diagonal reference line (AUC = 0.5) denotes the expected performance of a random classifier.

### Precision-Recall computation

Using the same ranked lists and threshold sweep in ROC computation, precision and recall at each threshold and average precision (AP) were computed as:

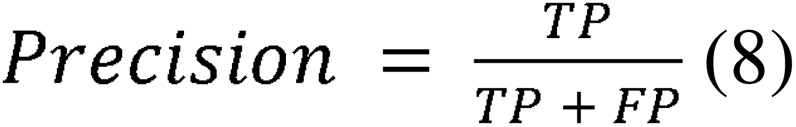

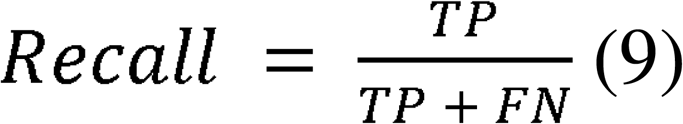

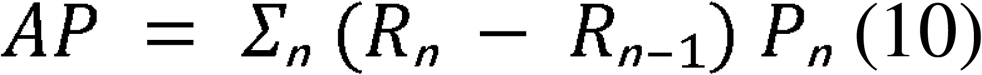

where P and R are the precision and recall at the n^th^ threshold. We also report precision and recall among the top k ranked compounds, with k set to the number of hits in each evaluated set (k = 324 for primary screen and 74 for validation screen). At this cutoff, precision@k equals recall@k (**Table S9**).

To provide a reference point that isolates the value of agentic reasoning, we implemented a fixed mean-aggregation baseline. This pipeline reads the same raw assay files supplied to the agent, harmonizes compound identifiers to ChEMBL IDs, and merges the assay results per compound on that identifier. Repeated measurements for a given compound were collapsed by a plain arithmetic mean; for dose-ranged assays, this averages across all tested concentrations, with no best-concentration selection, dose-response modeling, or toxicity filtering. Each assay readout was then standardized by min-max scaling and oriented so that higher values correspond to more hit-like behavior. The standardized readouts were averaged across the assays available for each compound to yield a composite score, and compounds were ranked by this score. The baseline was evaluated with the same ROC and precision-recall procedures applied to RepurAgent. The baseline was computed for both the primary screen (cell-painting morphology, CPE inhibition, and antibody-based infection rate) and the validation screen. The code is available in the Zenodo repository (**Data Availability**).

### Integrated services

“literature_search_pubmed” tool queries the NCBI LitSense 2.0 API^83^, a semantic search service developed and maintained by the National Center for Biotechnology Information. LitSense employs dense vector representations of biomedical text to enable semantic similarity search across the PubMed and PubMed Central corpora. The API was queried in "passages" mode, in which the index is searched at the sub-document level rather than at the level of full abstracts or articles.

Disease-specific biomedical KGs are generated via the KGG^50^ pipeline and represented as PyBEL objects. KG construction integrates multiple public resources, including:

- Open Targets Platform^61^: disease-associated targets and known drugs.

- ChEMBL^53^: molecule and mechanism of action lookups (via chembl-webresource-client) and canonical SMILES retrieval (via the ChEMBL REST API batch endpoint).

- UniProt^63^: protein annotation parsing from UniProt text records.

- Reactome^79^: pathway identifiers and browser links (including optional name-to-ID resolution).

- GWAS variants^62^: optional disease-associated variant retrieval via pandasgwas.

Physicochemical and biological annotation of the compound library was performed using the Chemical Annotator^52^ (v1.0), a unified chemical data annotation toolkit developed within the REMEDi4ALL framework. The tool accepts compound identifiers and performs exact-match queries against multiple public chemical databases, including ChEMBL^53^, UniChem^55^, PubChem^54^, and KEGG^56^.

For each compound, the annotator retrieves physicochemical properties together with biological activity data, including associated bioassay records, molecular targets, and quantitative activity measurements expressed as pChEMBL values. Annotation was restricted to high-confidence ChEMBL entries using a minimum confidence score threshold of 8, and only binding (B) and functional (F) assay types with a pChEMBL value ≥ 6 were retained, in line with the default recommended settings. Any additional compound metadata present in the input file was preserved in the output. The resulting annotated dataset was used as structured input for downstream computational analyses.

## Data availability

All agent execution traces, reasoning traces and analyses of agent outputs in this study are openly available in Zenodo at https://doi.org/10.5281/zenodo.24161441. Agent traces can additionally be browsed interactively through the deployed RepurAgent instance at https://repuragent.serve.scilifelab.se/.

## Code availability

The RepurAgent source code is available at https://github.com/pharmbio/repuragent-web. A standalone version for local deployment is available at https://github.com/pharmbio/repuragent.

## Acknowledgements

We thank Dr. Krister Wennerberg (Biotech Research and Innovation Centre, University of Copenhagen), Dr. Tom Erkers (Karolinska Institutet), Dr. Nadina Grosios (SMS-oncology), Dr. Jorrit Martijn Enserink (Oslo University Hospital), Dr. Sina Oppermann (Goethe-Universität Frankfurt), and Dr. Hong Yan (Goethe-Universität Frankfurt) for their expert evaluation of the AML repurposing candidates.

This work was supported by the Chemical Biology Consortium Sweden (CBCS), a national research infrastructure funded by the Swedish Research Council (dr.nr.2021-00179) and SciLifeLab. O.S. acknowledges funding from the Swedish Research Council (Grants 2024-04576, and 2024-03566), FORMAS (grants 2022-00940), Swedish Cancer Foundation (25 4914 Pj 01 H), and the Swedish strategic research programme eSSENCE. We acknowledge SciLifeLab Data Centre and the SciLifeLab Serve service for hosting the RepurAgent application.

This work was supported by the REMEDi4ALL project, which has received funding from the European Union’s Horizon Europe research and innovation programme under grant agreement No 101057442. Views and opinions expressed are those of the author(s) only and do not necessarily reflect those of the European Union, who cannot be held responsible for them.

## Author Contributions

D.L.H. and O.S. conceived and designed the study. D.L.H. developed the methodology, implemented the multi-agentic system, designed and performed the case studies. S.S. contributed to the methodology and analysis. F.B. contributed the Chemical Annotator tool. K.N. developed the RepurposeDrugs tool. R.K. and B.P. developed the knowledge graph tool. E.A., P.Ö., B.S.-L., J.C.P. and J.R. contributed to the design of the COVID-19 case study. A.D., Lars Schlotawa and Lina Schmidt designed and performed the MSD case study. D.L.H., O.S., J.C.P. and J.R. analysed and interpreted the results. T.A. contributed to interpretation of case studies and critically reviewed the work. O.S. supervised the project. D.L.H. and O.S. wrote the manuscript. All authors have read and approved the manuscript.

