## Supplementary Information for "Human-supervised Agentic AI for Hypothesis Generation and Experimental Assistance in Drug Repurposing"

**Table S1.** REMEDi4ALL SOPs integrated into the research agent

| **Name** | **Description** | **DOI** |
| --- | --- | --- |
| *In silico* drug repurposing tools and resources | Provides a workflow for identifying new uses for existing drugs using computational tools and databases. It covers target/pathway identification, compound selection (e.g., via ChEMBL), and prioritization based on physicochemical properties and safety data to support drug repurposing decisions. | <https://doi.org/10.5281/zenodo.15672626> |
| Drugs combination screening | Describes a dose-response matrix assay to evaluate interactions between drug combinations in cell-based systems. It includes experimental setup, cell seeding, treatment, viability measurement (luminescence), and data analysis to identify synergistic, additive, or antagonistic effects. | <https://doi.org/10.5281/zenodo.17206369> |
| A high-content screening assay to evaluate small molecule-induced phospholipidosis in cells | Outlines a standardized high-content imaging assay to evaluate small molecules–induced phospholipidosis in cell lines (e.g., U2OS, A549). | <https://doi.org/10.5281/zenodo.14872165> |
| Detection of Drug-Induced-Phospholipidosis | Describes a high-content screening (HCS) method to detect drug-induced phospholipidosis (DIPL) in VeroE6 cells using fluorescent staining and imaging. | <https://doi.org/10.5281/zenodo.13880870> |

**Table S2.** Tools in RepurAgent

| **Tool Name** | **Agent(s)** | **Description** |
| --- | --- | --- |
| python_executor | data_agent | Executing Python with restricted imports and persistent state between calls. |
| reset_python_state | data_agent | Clearing the persisted Python execution context. |
| literature_search_pubmed | planning_agent, research_agent | Querying LitSense API (v2.0) for literature passages. |
| protocol_search_sop | planning_agent, research_agent | Retrieving SOP/protocol content from the local SOP database. |
| annotate_chemicals | research_agent | Collecting drug annotations from public chemical databases (ChEMBL, UniChem, PubChem, and KEGG) based on exact match with query pattern. |
| search_disease_id | research_agent | Looking up disease EFO/MONDO IDs given disease names using OpenTarget API. |
| create_knowledge_graph | research_agent | Building disease-specific KG using real-time data from API calls. |
| extract_drugs_from_kg | research_agent | Extracting drugs and related targets/MoAs from the KG file |
| extract_mechanism_of_actions_from_kg | research_agent | Extracting mechanisms of action relationships from the KG file |
| extract_proteins_from_kg | research_agent | Extracting HGNC proteins/targets with druggability metadata from the KG file |
| extract_pathways_from_kg | research_agent | Extracting Reactome pathways linked to proteins/drugs from the KG file |
| getDrugsforProteins | research_agent | Retrieving drugs for provided proteins via OpenTargets |
| getDrugsforMechanisms | research_agent | Retrieving drugs associated with provided mechanisms of action. |
| getDrugsforPathways | research_agent | Retrieving drugs associated with provided pathways. |
| CYP3A4_classifier | prediction_agent | CPSign classifier predicting CYP3A4 inhibition. |
| hERG_classifier | prediction_agent | CPSign classifier for hERG inhibition risk. |
| AMES_classifier | prediction_agent | CPSign classifier for Ames mutagenicity. |
| PGP_classifier | prediction_agent | CPSign classifier for P-glycoprotein inhibition. |
| Solubility_regressor | prediction_agent | CPSign regression model providing solubility + prediction interval. |
| Lipophilicity_regressor (proxy) | prediction_agent | RDKit-based logP computation. |
| PAMPA_classifier | prediction_agent | CPSign classifier for PAMPA permeability assay. |
| BBB_classifier | prediction_agent | CPSign classifier estimating BBB penetration. |
| CYP2C19_classifier | prediction_agent | CPSign classifier for CYP2C19 inhibition. |
| CYP2D6_classifier | prediction_agent | CPSign classifier for CYP2D6 inhibition. |
| CYP1A2_classifier | prediction_agent | CPSign classifier for CYP1A2 inhibition. |
| CYP2C9_classifier | prediction_agent | CPSign classifier for CYP2C9 inhibition. |

**Table S3.** Configurations evaluated in the planning-agent ablation study

| **Configuration** | **Description** |
| --- | --- |
| P0-L0-E0 | Planning agent with vanilla LLM |
| P1-L0-E0 | Planning agent with protocol_search_sop tool |
| P0-L1-E0 | Planning agent with literature_search_pubmed tool |
| P0-L0-E1 | Planning agent with episodic_memory tool |
| P1-L1-E0 | Planning agent with protocol_search_sop and literature_search_pubmed tool |
| P1-L0-E1 | Planning agent with protocol_search_sop and episodic_memory tool |
| P0-L1-E1 | Planning agent with literature_search_pubmed and episodic_memory tool |
| P1-L1-E1 | Planning agent with literature_search_pubmed, protocol_search_sop, and episodic_memory tool |

**Table S4.** Scoring criteria for planning agent evaluation

| **Dimension** | **Guiding question** | **Score 1** | **Score 3** | **Score 5** |
| --- | --- | --- | --- | --- |
| **Structure & completeness** | *Is the plan coherent, ordered, and cover the task end-to-end?* | No order; major phases missing or jumbled | Recognizable structure but skips one or more necessary steps, or ordering inconsistent | Phases complete, sequenced correctly, clearly connected from problem framing to final output |
| **Task alignment** | *How well does it match the reference workflow?* | Addresses a different problem or ignores core workflow requirements | Covers main objective but misses/misframes specific steps or constraints | Every step maps cleanly to reference workflow; scope and framing fully consistent |
| **Executability** | *Are steps concrete enough to execute with realistic handoffs and outputs?* | Vague actions ('analyze data') with no named tools, methods, or outputs | Names outputs but leaves tool choice or handoff conditions underspecified | Each step names a tool/method, a concrete output artifact, and the condition for passing to the next step |
| **Scientific rigor** | *Sound scientific reasoning for drug repurposing?* | Scientifically unsound: wrong assumptions, inappropriate methods, missing validation logic | Reasonable overall but has methodological gap or unjustified assumption | Reflects best practice: appropriate rationale, suitable prioritization, realistic validation |
| **Context use** | *Does the plan use the enabled/provided context appropriately and productively?* | Context ignored, or hallucinated sources/evidence not provided | Context acknowledged but used inconsistently | Context tightly integrated; specifics inform decisions and are cited accurately; nothing unsupported invented |
| **Specificity** | *Specific and non-generic rather than boilerplate?* | Fully generic; could describe any repurposing project; no disease/target/dataset specifics | Some specific elements, but key sections read as templated filler | Highly specific: disease, target rationale, data sources, thresholds, methods named and tailored |

**Table S5.** Human plan revisions in each case study session.

| **Session** | **Rounds of revision** | **Scientist feedbacks** | **Reflected in execution/outputs** |
| --- | --- | --- | --- |
| AML | 1 | - Step 2 should merge previous files before filter out non-small-molecular drugs.  - Step 4 should prioritize mechanistic scores over ADMET  - Step 4 should output a full ranked list of drugs before extracting the top 20.  - Step 5 should be incorporated into step 4 as the last sub-task. Combine these steps | - master_dataset.csv and master_compounds_dedup_by_inchikey.csv exported  - overall_score = 0.85 × mechanistic_norm + 0.15 × (ADMET bonus − 0.15 × ADMET penalty)  - full_ranked_list.csv (397 compounds) before top20_for_experimental_testing.csv |
| COVID-19 primary (unconstrained) | 1 | - Step 5 should search for SOP and literature to suggest next step | - Research Agent ran 1 SOP search and 3 literature searches to propose validation experiments; normalization, weighting and ranking were unchanged and agent-derived |
| COVID-19 primary (constrained) | 2 | - Round 1: Combine every step into one single step handled by a data agent. For the identifying confounding, analyze why one compound works in one assay but not others.  - Round 2: Add a step at the end call Report and documenting, handled by Report Agent | - discrepancy_explanations_top10pct_discordance.csv |
| COVID-19 validation | 1 | - Combine steps 1-5 into a single step. This step should also document what it does into a md file | - Single combined analysis step; analysis_documentation.md; ranking scheme agent-derived |
| COVID-19 final selection | 2 | - Round 1: Before integrate data, try to read and understand it first. The assay performance should consider all available assay readouts (morphology score, infection rate, cytopathicity inhibition, etc). When developing a ranking system, treat all factors as equal. The markdown files for each of top 5 drugs should be well formatted with tables and figures if needed. Should use the analysed data as evidence and interpret them.    - Round 2: Combine steps 2-10 into a single step. | - ranking_report.md: 'Equal-weight ranking system (8 factors)', composite = mean of eight 0–1 sub-scores  - Research Agent annotations used as inputs; top-5 dossiers with tables and figures.  - After round 2 the plan was consolidated into annotation retrieval + one Data Agent step |
| MSD | 1 | - Combine step 1 and 2 together / Combine step 3 and 4 together | - Two-step plan (Research Agent: disease profile + library annotation; Data Agent: screening + tiering) before reporting; therapeutic directions and tier criteria derived by the agent |

**Table S6.** Biological pathways shared by RepurAgent's and Google Co-Scientist's suggested drugs for AML

| **Group** | **Pathways** |
| --- | --- |
| FLT3 signaling | FLT3 Signaling; FLT3 Signaling In Disease; FLT3 Signaling By CBL Mutants; FLT3 Signaling Thru SRC Family Kinases; Signaling By FLT3 ITD And TKD Mutants; Negative Regulation Of FLT3; STAT5 Activation Downstream Of FLT3 ITD Mutants |
| MAPK / RAS / RAF | MAPK Family Signaling Cascades; MAPK1/MAPK3 Signaling; MAP Kinase Activation; MAPK1 (ERK2) Activation; MAPK3 (ERK1) Activation; Oncogenic MAPK Signaling; Negative Feedback Regulation Of MAPK Pathway; RAF/MAP Kinase Cascade; RAF Activation; RAF-Independent MAPK1/3 Activation; Signaling By BRAF And RAF1 Fusions; Signaling By High-Kinase Activity BRAF Mutants; Signaling By RAF1 Mutants; Paradoxical Activation Of RAF Signaling By Kinase Inactive BRAF; Prolonged ERK Activation Events; Signaling To ERKs |
| PI3K / AKT | PI3K Cascade; PI3K/AKT Signaling In Cancer; Constitutive Signaling By Aberrant PI3K In Cancer; PIP3 Activates AKT Signaling; Negative Regulation Of PI3K/AKT Network; PI5P, PP2A And IER3 Regulate PI3K/AKT Signaling; Intracellular Signaling By Second Messengers |
| JAK-STAT | STAT5 Activation; Interferon Gamma Signaling; Interferon Signaling; Regulation Of IFNG Signaling; Growth Hormone Receptor Signaling; Prolactin Receptor Signaling |
| Cytokine & hematopoietic growth factor signaling | Cytokine Signaling In Immune System; Signaling By Interleukins; Interleukin-3, Interleukin-5 And GM-CSF Signaling; Interleukin-6 Signaling; Interleukin-6 Family Signaling; IL-6-Type Cytokine Receptor Ligand Interactions; Interleukin Receptor SHC Signaling; Signaling By Erythropoietin; Erythropoietin Activates PI3K; Erythropoietin Activates PLCG; Erythropoietin Activates RAS; Erythropoietin Activates STAT5; Signaling By CSF3 (G-CSF); Inactivation Of CSF3 (G-CSF) Signaling; Factors Involved In Megakaryocyte Development And Platelet Production; Interleukin-2 Family Signaling; Interleukin-4 And Interleukin-13 Signaling; Interleukin-12 Family Signaling; Interleukin-12 Signaling; Interleukin-17 Signaling; Interleukin-20 Family Signaling; Interleukin-23 Signaling; Interleukin-27 Signaling; Interleukin-35 Signaling; Interleukin-1 Family Signaling; Interleukin-1 Signaling |
| RTK signaling (KIT, IGF1R, insulin, NTRK) | Signaling By Receptor Tyrosine Kinases; Signaling By KIT In Disease; Signaling By SCF-KIT; IGF1R Signaling Cascade; IRS-Related Events Triggered By IGF1R; Signaling By Type 1 IGF1 Receptor; Insulin Receptor Signaling Cascade; Signaling By Insulin Receptor; IRS-Mediated Signaling; Signaling By NTRKs; Signaling By NTRK1 (TRKA); Frs2-Mediated Activation; Signaling By Leptin |
| Cell cycle | Cyclin D Associated Events In G1; Mitotic G1 Phase And G1/S Transition |
| Innate immunity / TLR signaling | Immune System; Toll-Like Receptor Cascades; Toll Like Receptor 3 (TLR3) Cascade; Toll Like Receptor 4 (TLR4) Cascade; Toll Like Receptor 7/8 (TLR7/8) Cascade; Toll Like Receptor 9 (TLR9) Cascade; MyD88 Cascade Initiated On Plasma Membrane; MyD88 Dependent Cascade Initiated On Endosome; MyD88-Independent TLR4 Cascade; MyD88:MAL(TIRAP) Cascade Initiated On Plasma Membrane; TRAF6 Mediated Induction Of NFkB And MAP Kinases Upon TLR7/8 Or 9 Activation |
| Chromatin modification & epigenetics | Chromatin Modifying Enzymes; RMTs Methylate Histone Arginines |
| Nervous system / axon guidance | Nervous System Development; Axon Guidance; Developmental Biology; L1CAM Interactions; Signal Transduction By L1 |
| Disease / broad signal transduction | Signal Transduction; Disease; Diseases Of Signal Transduction By Growth Factor Receptors And Second Messengers; Infectious Disease; SARS-CoV Infections; Potential Therapeutics For SARS |

**Table S7.** Holm-adjusted p-values for RepurAgent versus vanilla LLMs/random baseline on pathway-overlap metrics with Google Co-Scientist, with gradually excluding broad Reactome terms.

| **Metric** | **RepurAgent vs** | **Broad terms excluded** | | | | | | |
| --- | --- | --- | --- | --- | --- | --- | --- | --- |
|  |  | **None (106)** | **> 500 (98)** | **> 300 (92)** | **> 200 (88)** | **> 100 (68)** | **> 50 (53)** | **> 20 (27)** |
| **Recall** | GPT-4o | 2.6×10⁻⁴*** | 2.6×10⁻⁴*** | 2.6×10⁻⁴*** | 2.6×10⁻⁴*** | 2.6×10⁻⁴*** | 2.6×10⁻⁴*** | 2.6×10⁻⁴*** |
|  | GPT-4o-mini | 0.005** | 0.005** | 0.005** | 0.003** | 0.003** | 6.1×10⁻⁴*** | 2.6×10⁻⁴*** |
|  | GPT-5 | 0.016* | 0.008** | 0.008** | 0.008** | 0.003** | 0.002** | 7.1×10⁻⁴*** |
|  | GPT-5-mini | 2.6×10⁻⁴*** | 2.6×10⁻⁴*** | 2.6×10⁻⁴*** | 2.6×10⁻⁴*** | 2.6×10⁻⁴*** | 2.6×10⁻⁴*** | 2.6×10⁻⁴*** |
|  | GPT-5.1 | 3.0×10⁻⁴*** | 2.8×10⁻⁴*** | 2.8×10⁻⁴*** | 3.0×10⁻⁴*** | 2.6×10⁻⁴*** | 2.6×10⁻⁴*** | 2.6×10⁻⁴*** |
|  | GPT-5.2 | 2.6×10⁻⁴*** | 2.6×10⁻⁴*** | 2.6×10⁻⁴*** | 2.6×10⁻⁴*** | 2.6×10⁻⁴*** | 2.6×10⁻⁴*** | 2.6×10⁻⁴*** |
|  | Random: ChEMBL cancer drugs | 2.6×10⁻⁴*** | 2.6×10⁻⁴*** | 2.6×10⁻⁴*** | 2.6×10⁻⁴*** | 2.6×10⁻⁴*** | 2.6×10⁻⁴*** | 2.6×10⁻⁴*** |
|  | Random: initial drug space | 0.009** | 0.006** | 0.005** | 0.004** | 0.003** | 0.003** | 0.003** |
| **Precision** | GPT-4o | 0.016* | 0.006** | 0.005** | 0.003** | 0.001** | 0.003** | 0.004** |
|  | GPT-4o-mini | 0.002** | 2.6×10⁻⁴*** | 2.6×10⁻⁴*** | 2.6×10⁻⁴*** | 2.6×10⁻⁴*** | 2.6×10⁻⁴*** | 2.6×10⁻⁴*** |
|  | GPT-5 | 2.6×10⁻⁴*** | 2.6×10⁻⁴*** | 2.6×10⁻⁴*** | 3.0×10⁻⁴*** | 2.6×10⁻⁴*** | 2.6×10⁻⁴*** | 2.6×10⁻⁴*** |
|  | GPT-5-mini | 0.001** | 0.001** | 9.1×10⁻⁴*** | 0.001** | 2.6×10⁻⁴*** | 2.6×10⁻⁴*** | 2.6×10⁻⁴*** |
|  | GPT-5.1 | 0.001** | 0.001** | 0.001** | 0.001** | 2.9×10⁻⁴*** | 6.1×10⁻⁴*** | 2.6×10⁻⁴*** |
|  | GPT-5.2 | 2.6×10⁻⁴*** | 2.6×10⁻⁴*** | 2.6×10⁻⁴*** | 2.6×10⁻⁴*** | 2.6×10⁻⁴*** | 2.6×10⁻⁴*** | 2.6×10⁻⁴*** |
|  | Random: ChEMBL cancer drugs | 0.009** | 0.006** | 0.005** | 0.002** | 2.9×10⁻⁴*** | 2.6×10⁻⁴*** | 2.6×10⁻⁴*** |
|  | Random: initial drug space | 0.684 (n.s.) | 0.565 (n.s.) | 0.529 (n.s.) | 0.529 (n.s.) | 0.529 (n.s.) | 0.684 (n.s.) | 0.218 (n.s.) |
| **Jaccard index** | GPT-4o | 0.002** | 0.001** | 0.001** | 9.1×10⁻⁴*** | 0.001** | 0.002** | 0.004** |
|  | GPT-4o-mini | 2.6×10⁻⁴*** | 2.6×10⁻⁴*** | 2.6×10⁻⁴*** | 2.6×10⁻⁴*** | 2.6×10⁻⁴*** | 2.6×10⁻⁴*** | 2.6×10⁻⁴*** |
|  | GPT-5 | 3.0×10⁻⁴*** | 5.2×10⁻⁴*** | 9.1×10⁻⁴*** | 9.1×10⁻⁴*** | 2.6×10⁻⁴*** | 2.6×10⁻⁴*** | 2.6×10⁻⁴*** |
|  | GPT-5-mini | 2.6×10⁻⁴*** | 2.6×10⁻⁴*** | 2.6×10⁻⁴*** | 2.6×10⁻⁴*** | 2.6×10⁻⁴*** | 2.6×10⁻⁴*** | 2.6×10⁻⁴*** |
|  | GPT-5.1 | 9.1×10⁻⁴*** | 7.1×10⁻⁴*** | 9.1×10⁻⁴*** | 9.1×10⁻⁴*** | 2.6×10⁻⁴*** | 2.6×10⁻⁴*** | 2.6×10⁻⁴*** |
|  | GPT-5.2 | 2.6×10⁻⁴*** | 2.6×10⁻⁴*** | 2.6×10⁻⁴*** | 2.6×10⁻⁴*** | 2.6×10⁻⁴*** | 2.6×10⁻⁴*** | 2.6×10⁻⁴*** |
|  | Random: ChEMBL cancer drugs | 2.6×10⁻⁴*** | 2.6×10⁻⁴*** | 2.6×10⁻⁴*** | 2.6×10⁻⁴*** | 2.6×10⁻⁴*** | 2.6×10⁻⁴*** | 2.6×10⁻⁴*** |
|  | Random: initial drug space | 0.037* | 0.044* | 0.037* | 0.037* | 0.105 (n.s.) | 0.210 (n.s.) | 0.087 (n.s.) |

Each cell contains the Holm-adjusted p-value of a two-sided Mann–Whitney U test comparing RepurAgent with the vanilla LLMs/random baselines over 10 independent runs each; the adjustment is applied within each exclusion rule over 24 tests (3 metrics × 8 comparators). At each threshold N, Reactome pathways that have gene sets larger than N genes were removed from both the Co-Scientist reference set and every arm's pathway set before the metrics were recomputed. * p < 0.05, ** p < 0.01, *** p < 0.001; n.s., not significant (shaded).

**Table S8.** Rubric for the expert evaluation in the AML case study

| **Criterion** | **What it measures** | **Score 1** | **Score 3** | **Score 5** |
| --- | --- | --- | --- | --- |
| **A1. Mechanistic plausibility for AML** | Strength of support that the drug's known pharmacology produces an anti-leukaemic effect. Assesses the mechanism (target/pathway/MoA), not whether the drug itself has previously been studied in AML. | No coherent causal path from the mechanism to an anti-leukaemic effect, or the proposed link is contradicted by known AML biology. | The mechanism is a biologically reasonable route to an anti-leukaemic effect, but the link rests mainly on inference or analogy (structural reasoning, in silico signal, extrapolation from a related setting) without direct experimental or literature support. | Direct experimental or strong biological demonstration in prior literature that the mechanism produces an anti-leukaemic effect. |
| **A2. Specificity of target/pathway rationale** | How AML-specific the drug's target/pathway/MoA is, on a spectrum from pan-cancer to AML-defining. | Only broad, pan-cancer processes (apoptosis, proliferation, generic cytotoxicity), with nothing specifically relevant to leukaemia. | A mix of general and some AML-relevant mechanisms. | A precise, AML-defining node: a specific molecular target, driver lesion or dysregulated pathway (e.g. FLT3-ITD, IDH1/2, mutant NPM1, menin–KMT2A, BCL-2 dependence, differentiation block). |
| **A3. Novelty for repurposing** | How much prior attention the drug–AML hypothesis has already received. | Established or approved in AML. | Previously proposed for AML: one or more prior studies, trials, screens or reviews have raised the drug–AML link. | No prior AML-specific investigation known; the connection appears new for this disease. |
| **A4. Druggability / developability for AML** | How practically feasible it is to advance the drug in AML. | A prohibitive liability makes development unrealistic: unreachable therapeutic window, unacceptable added myelosuppression, or a severe unavoidable drug–drug interaction with standard AML care. | Feasible, but with meaningful caveats to manage (dose or schedule optimisation, monitoring, formulation work, manageable toxicity). | Readily developable: favourable, well-understood exposure with a credible therapeutic window, a safety profile compatible with the elderly, myelosuppressed, polypharmacy AML setting, and a low barrier to clinical testing. |

**Table S9.** Precision and recall among the top-ranked compounds in the COVID-19 case study.

| **Stage** | **Methods** | **k** | **precision@k = recall@k** |
| --- | --- | --- | --- |
| Primary screen | RepurAgent (constrained) | 324 | 1.0 |
| Primary screen | Repuragent assistant | 324 | 0.816 |
| Primary screen | Naive baseline | 324 | 0.642 |
| Validation screen | Repuragent assistant | 74 | 0.699 |
| Validation screen | Naive baseline | 74 | 0.548 |

Compounds were ranked by each method, and the top k compounds were taken as predicted hits, with k equal to the number of expert-selected hits in the evaluated set. At this cutoff, precision@k equals recall@k.

**Table S10.** Confounder flags raised by RepurAgent when analyzing assay readouts

| **Stage** | **Confounder flag** | **Description** | **Number of compounds** |
| --- | --- | --- | --- |
| **Primary Screen** | flag_ab_toxicity | strong AB activity and high cytotoxicity (low cell viability). Possibly reducing infection may be driven by cell loss. | 242 |
|  | flag_cpe_toxicity | strong CPE rescue and high toxicity (low cell viability). Possible assay interference / non-specific effects; also indicates narrow therapeutic window. | 595 |
|  | flag_cp_only | strong CP morphology rescue but no AB/CPE support. Morphology rescue may be unrelated to viral suppression. | 24 |
|  | flag_ab_no_cpe | AB strong but mean CPE negative → inconsistent antiviral and cytotoxicity readouts; warrants follow-up. | 241 |
| **Validation screen** | morphology_rescue_  without_infection_reduction | Morphology_score* ≥ 0.5 but infection rate stayed high. | 9 |

* Morphology score: distance to a healthy (non-infected) control, quantified from morphological features

**Table S11.** ADMET model performance on external test sets

| **Classification Models** | | | | | | | |
| --- | --- | --- | --- | --- | --- | --- | --- |
| **Model** | **Confidence** | **Accuracy** | **Accuracy (0)** | **Accuracy (1)** | **Proportion empty-label prediction sets** | **Proportion multi-label prediction sets** | **Proportion single-label prediction sets** |
| BBB | 0.8 | 0.8 | 0.875 | 0.777 | 0 | 0.0173 | 0.983 |
| AMES | 0.8 | 0.786 | 0.777 | 0.793 | 0.0153 | 0.000695 | 0.984 |
| CYP1A2 | 0.8 | 0.822 | 0.828 | 0.816 | 0.000398 | 0.0111 | 0.988 |
| CYP2C9 | 0.8 | 0.811 | 0.814 | 0.806 | 0 | 0.123 | 0.877 |
| CYP2C19 | 0.8 | 0.807 | 0.808 | 0.807 | 0 | 0.0774 | 0.923 |
| CYP2D6 | 0.8 | 0.805 | 0.802 | 0.815 | 0 | 0.163 | 0.837 |
| CYP3A4 | 0.8 | 0.804 | 0.801 | 0.808 | 0 | 0.104 | 0.896 |
| hERG | 0.8 | 0.805 | 0.805 | 0.818 | 0 | 0.0698 | 0.93 |
| PAMPA | 0.8 | 0.789 | 0.78 | 0.79 | 0 | 0.305 | 0.695 |
| PGP | 0.8 | 0.816 | 0.763 | 0.862 | 0.0533 | 0 | 0.947 |
| **Regression models** | | | | | | | |
| **Model** | **Confidence** | **MAE** | **RMSE** | **Median interval width** | | **Accuracy** | |
| Solubility | 0.71 | 0.859 | 1.45 | 1.9 | | 0.716 | |

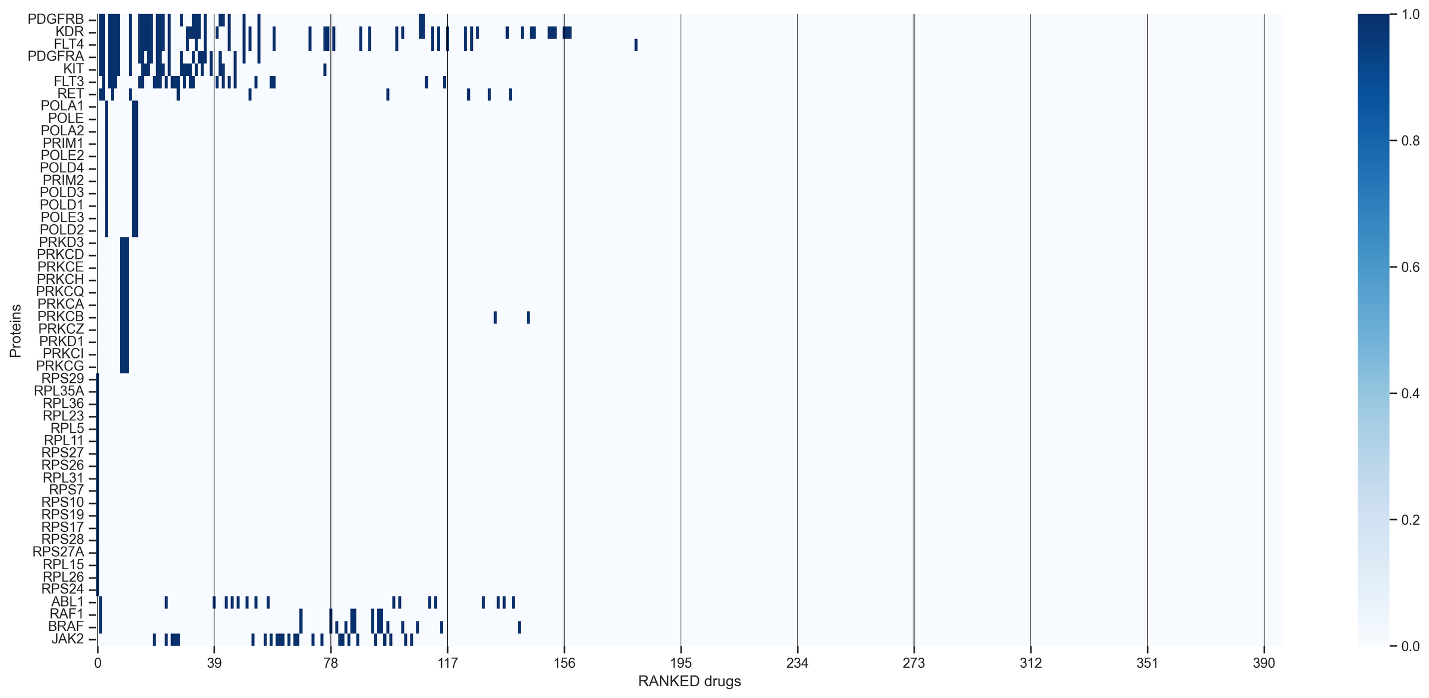

**Figure S1. Drug–protein heatmap across all ranked compounds.** Each row represents a distinct protein and each column a ranked drug candidate. The sparse pattern reflects the selectivity of top-ranked compounds for specific proteins, with enrichment in kinases, platelet-derived growth factor receptors, ribosomal proteins, and DNA polymerases.

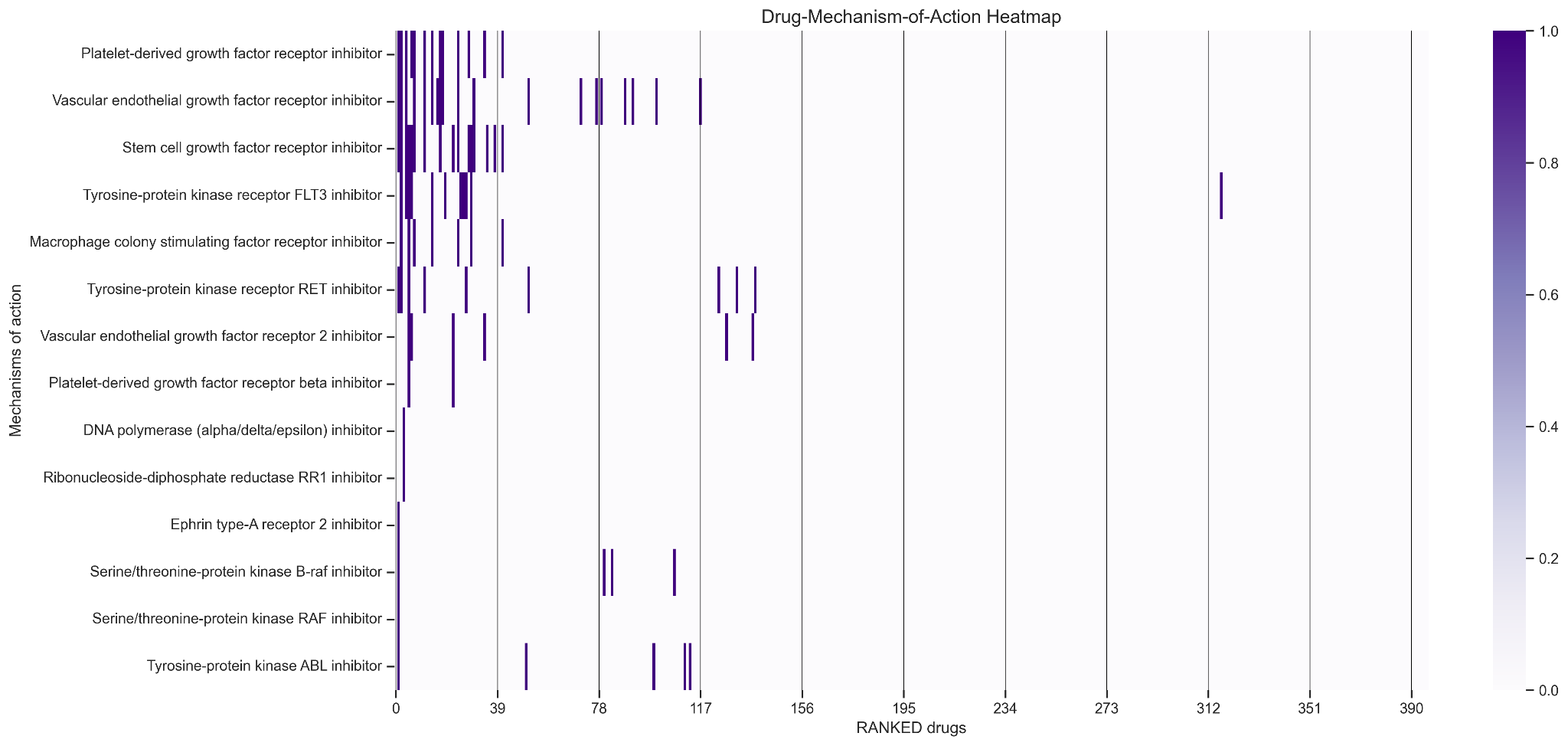

**Figure S2.** **Drug–mechanism-of-action heatmap across all ranked compounds.** Each row represents a distinct mechanism of action and each column a ranked drug candidate. The sparse pattern reflects the selectivity of top-ranked compounds for specific mechanistic classes, with notable enrichment in receptor tyrosine kinase and growth factor receptor inhibitor categories among the highest-ranked candidates.

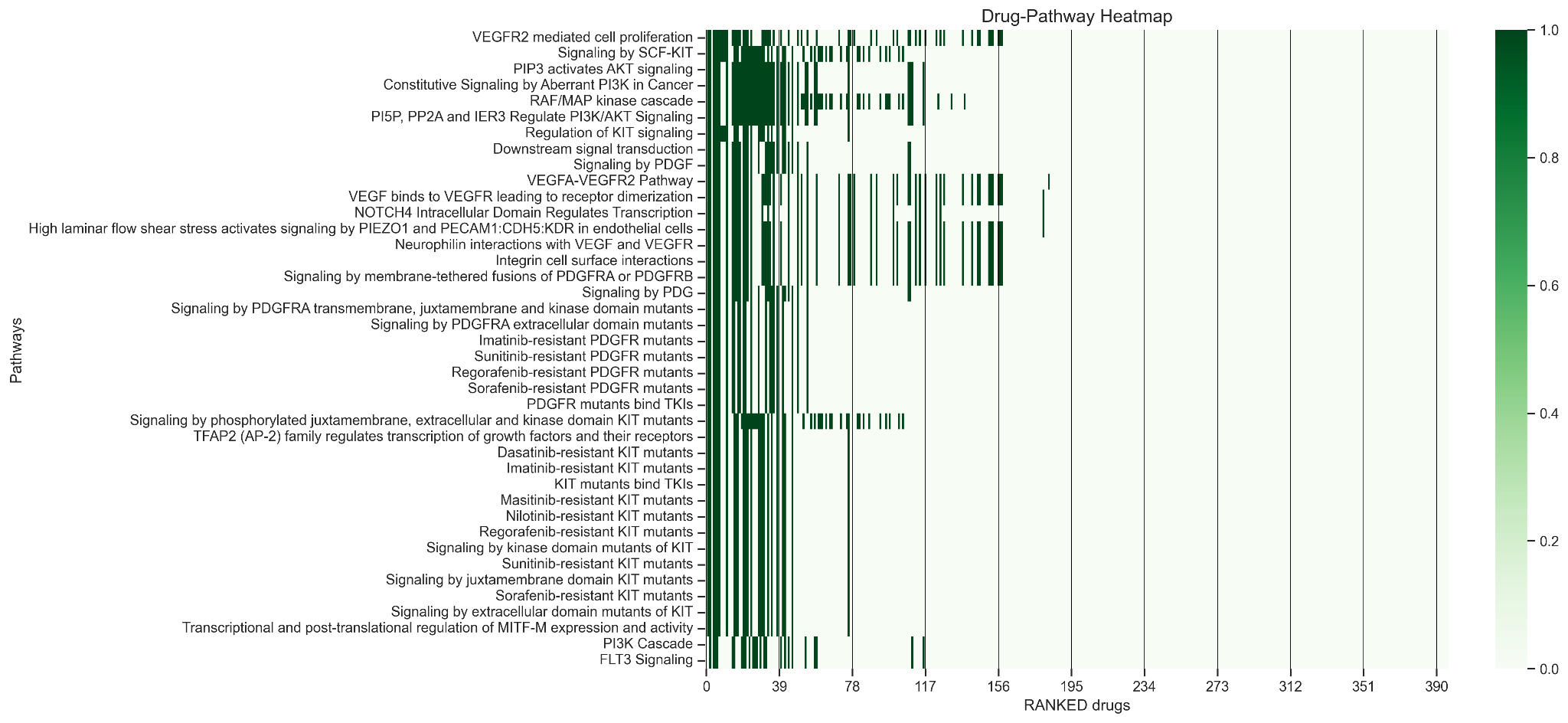

**Figure S3. Drug–pathway heatmap across all ranked compounds.** Each row represents a biological signaling pathway and each column a ranked drug candidate. Prominent pathway clusters include VEGFR-mediated cell proliferation, AKT and PDGFR signaling, and resistance-associated mutant pathways, which indicates both mechanistic diversity and convergent pathway targeting among the prioritized drug candidates

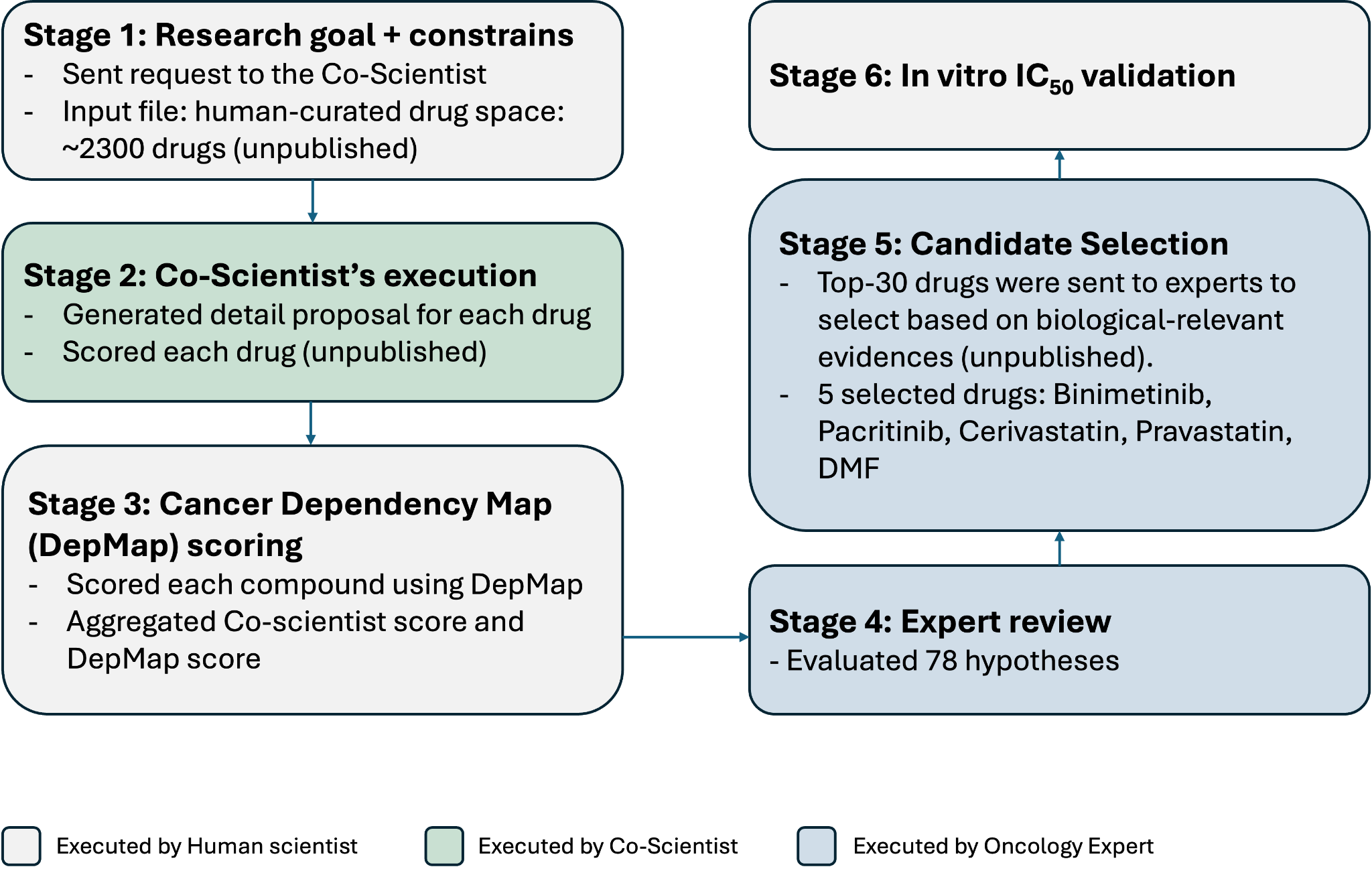

**Figure S4. Google Co-Scientist workflow for identifying drug repurposing candidates for acute myeloid leukemia (AML).** This workflow integrates contributions from a human scientist, the AI Co-Scientist, and external oncology experts. The Co-Scientist's outputs serve as one component within a broader expert-driven process, resulting in the selection of five candidates for in vitro IC₅₀ validation. Stages marked "unpublished" indicate data not reported in the publication.

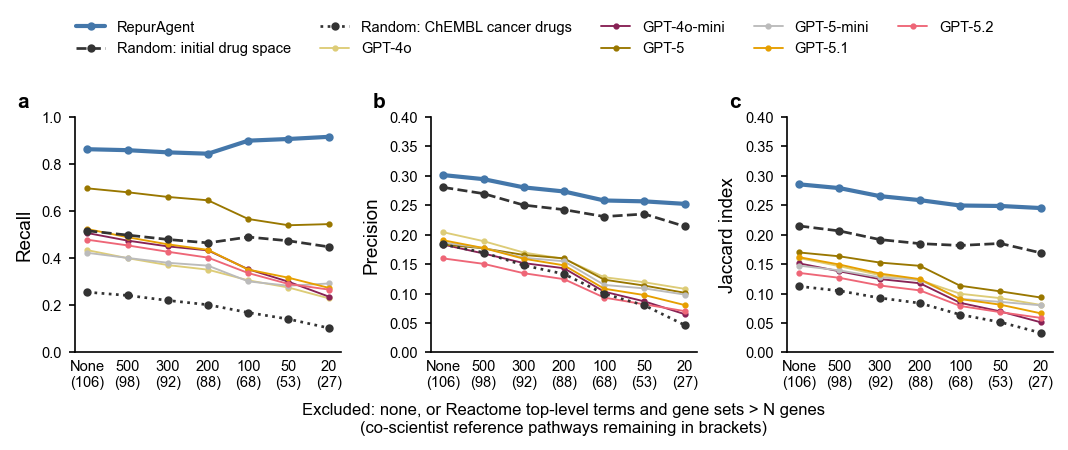

**Figure S5. Sensitivity of the pathway overlaps with Google Co-Scientist to the exclusion of broad Reactome terms.** (a) Recall, (b) precision and (c) Jaccard index of each arm's pathway set against the Co-Scientist reference set, as broad terms are gradually removed. At each threshold N, pathways with gene sets larger than N genes were removed from both the Co-Scientist reference set and every arm's pathway set before the metrics were recomputed. The numbers in brackets give the Co-Scientist reference pathways remaining (106 without exclusion, 27 at N = 20). Lines show the mean over 10 independent runs per arm. Under every threshold, RepurAgent remained significantly above the random cancer-drug baseline and all vanilla LLMs on all three metrics (two-sided Mann–Whitney U, p ≤ 0.016).

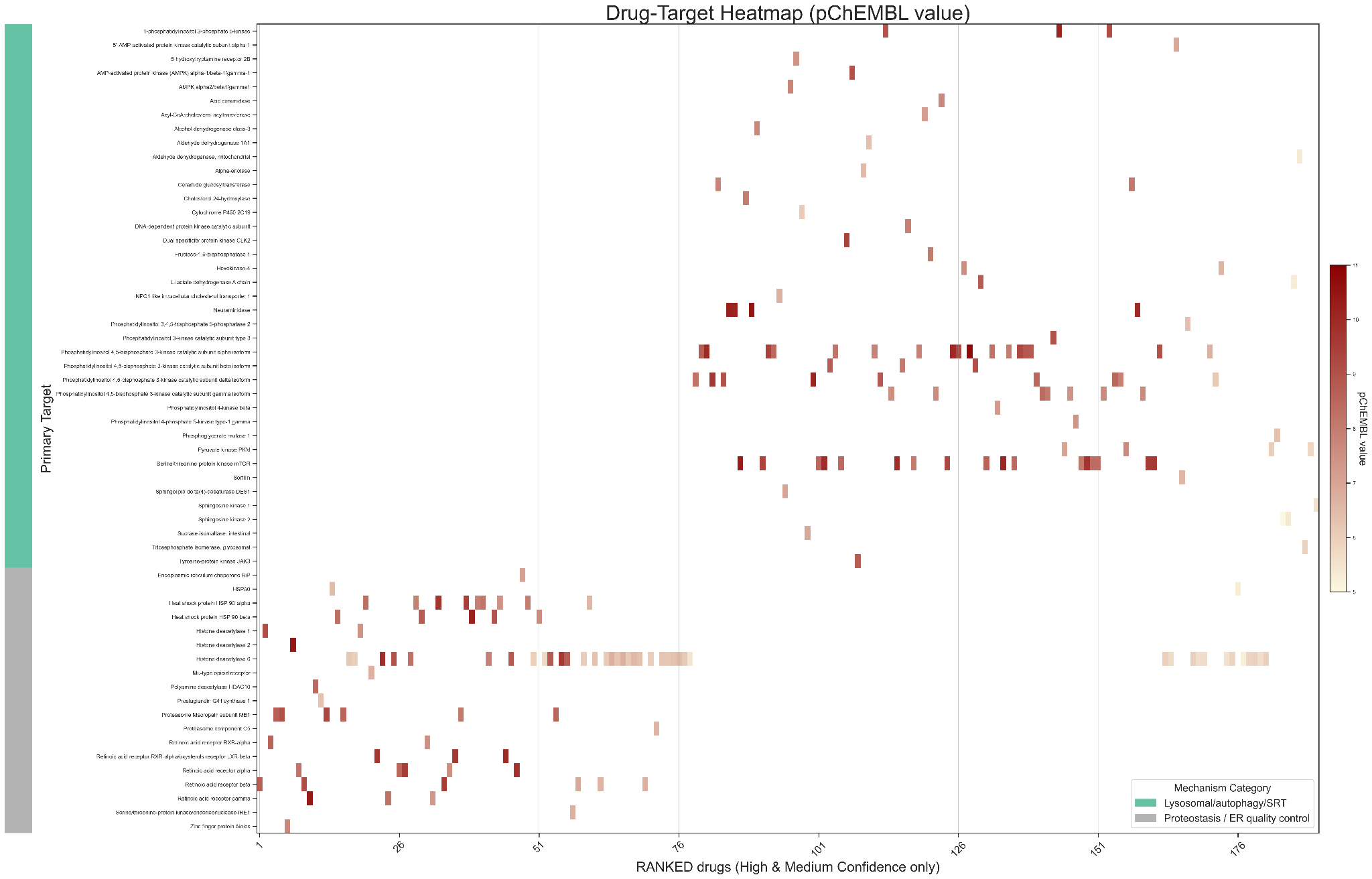

**Figure S6.** Drug-target heatmap illustrating the pChEMBL binding affinity values between high and medium confidence compounds (x-axis) and their annotated protein targets (y-axis). Cell color intensity reflects pChEMBL value, with darker red indicating stronger predicted binding affinity. Targets are grouped by mechanism of action category (lysosomal/autophagy/SRT shown in teal; proteostasis/ER quality control in green), enabling visualization of the pharmacological landscape and target coverage across the prioritized compound set.

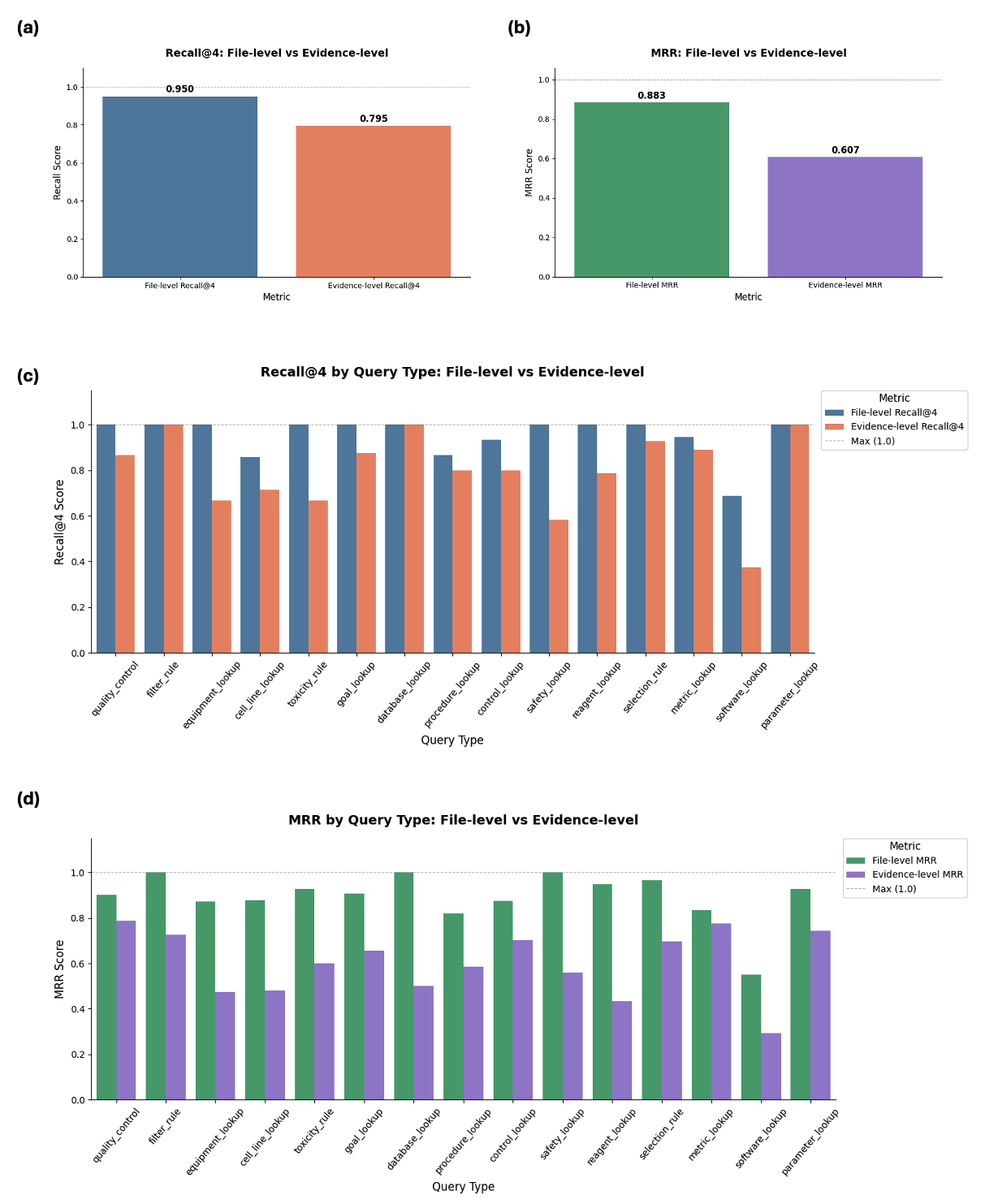

**Figure S7. Retrieval performance of the RAG system across evaluation levels and query types.** (a) Overall Recall@4 at file level and evidence level, measured over the top-4 retrieved chunks (k = 4). (b) Overall Mean Reciprocal Rank (MRR) at file level and evidence level, computed across all retrieved chunks to assess result prioritisation. (c) Recall@4 stratified by query type, comparing file-level and evidence-level scores across 15 query categories. (d) MRR stratified by query type, comparing file-level and evidence-level prioritisation performance across the same 15 categories. Dashed lines indicate the maximum achievable score (1.0). All metrics were evaluated on a stratified benchmark of 200 queries.

### **Note 1. Technical details of episodic memory design**

**Trigger.** Episodic memory was designed to personalise planning: it lets a user keep the planning experience they want RepurAgent to reuse. Nothing is saved automatically, instead, at the end of a session, the user can press “Extract Learning” in the interface when they judge the session worth reusing.

**Extraction and storage.** When extraction is triggered, the conversation of the current session is loaded from the short-term memory store and an extraction model (gpt-4o-mini) summarizes it into one structured output (one episode) with four fields:

- Task: the user's request.
- Initial decomposition: the step sequence the planning agent first proposed.
- Final decomposition: the step sequence after all of the user's revisions.
- Notes: lessons from the session that could improve a future attempt.

Episodes are stored in a ChromaDB (v0.6.3) collection and embedded with ChromaDB's default sentence encoder (all-MiniLM-L6-v2, 384 dimensions), using L2 distance.

**Retrieval.** For a new request, the two nearest episodes are retrieved and inserted into the planning agent's system prompt as worked examples (dynamic few-shot learning). Each example shows the task, the final decomposition and the notes, in the same format the planner is asked to produce. The initial plan is not shown to avoid confusion. Episodic memory acts on the planning agent directly; the other agents are influenced only through the plan.
